# One probe fits all: a highly customizable modular RNA *in situ* hybridization platform expanding the application of SABER DNA probes

**DOI:** 10.1101/2024.05.22.595454

**Authors:** Kirill Ustyantsev, Mattia Stranges, Filippo Giovanni Volpe, Stijn Mouton, Eugene Berezikov

## Abstract

*In situ* hybridization (ISH) of RNA is a key method to visualize gene expression patterns in complex biological samples. The technique is indispensable for biological research related to e.g. development, disease, gene function, and the validation of novel cell types identified by single-cell sequencing methods. Especially in non-mammalian models lacking accessibility to a broad spectrum of antibodies, ISH remains a major research tool. Diverse available ISH protocols require different custom hybridization probe types, design, and/or proprietary signal detection chemistry. This makes it hard to navigate for a beginner and increases the research costs when multiple methods need to be applied. Here, we describe OneSABER – a unified open platform connecting commonly used canonical and recently developed single- and multiplex, colorimetric, and fluorescent ISH approaches. This platform uses a single type of ISH DNA probes adapted from the signal amplification by exchange reaction (SABER) method. We demonstrate applications of the proposed ISH framework in whole-mount samples of the regenerative flatworm *Macrostomum lignano*, advancing this animal as a powerful model for stem cell and regeneration research.

## INTRODUCTION

Dynamic changes of spatiotemporal gene expression in different cell types, tissues, and organs reflect the complex molecular regulation and functioning taking place in an organism in response to internal and external stimuli. Studying these dynamics is crucial for understanding the roles of specific genes in key biological processes, such as embryonic development, homeostasis, and regeneration, in both normal and disease conditions. RNA *in situ* hybridization (ISH) and immunohistochemistry methods have allowed researchers to directly detect, measure, and visualize transcriptional and post-translational expression patterns of genes within native anatomical and histological contexts, connecting gene expression to the source cell (Hemmati-Brivanlou et al., 1990; Jowett and Lettice, 1994; Tautz and Pfeifle, 1989; Toledano et al., 2012; Young et al., 2020). Due to paucity or even complete absence of commercial antibodies for most non-mammalian species (Hadwiger et al., 2010; Peter et al., 2017), ISH remains the main research tool to study gene expression patterns in non-canonical models. Furthermore, rapid expansion and broad accessibility of next-generation single-cell sequencing technologies enabled the discovery of potential novel cell types and their marker genes, which requires experimental validation by *in situ* methods (Chari et al., 2021; Fincher et al., 2018; Hulett et al., 2023).

For decades, enzyme-catalyzed reporter deposition assays such as alkaline phosphatase (AP) colorimetric ISH and horseradish peroxidase tyramide signal amplification (TSA) fluorescent ISH (FISH) have been the gold standards in achieving reliable high signal-over-noise RNA and protein assays in whole-mount preparations (Bobrow et al., 1989; Jowett and Lettice, 1994; Luehrsen et al., 2000; Schmidt et al., 1997; Tautz and Pfeifle, 1989; Young et al., 2020). To date, many different open-access and commercial (F)ISH approaches have been developed, focusing on optimizing the methods’ sensitivity (signal amplification), specificity (low off-target signal), resolution (down to subcellular localization), and protocol implementation time, while allowing multiplexing - simultaneous visualization of multiple target molecules in the same sample (Dardani et al., 2022; Goh et al., 2020; Kishi et al., 2019; Schulte et al., 2024; Tao et al., 2023; Wang and Guo, 2021; Wang et al., 2012; Wu et al., 2022; Xia et al., 2019; Young et al., 2020). Excelling in all these aspects of ISH is a big challenge when working with complex, thick, and highly autofluorescent whole-mount samples. This leaves many of the mostly multiplexing-focused tools unpractical beyond the levels of cell cultures or thin tissue sections (Schulte et al., 2024). Therefore, there is still room for the canonical AP-based and TSA FISH methods, despite their limited multiplexing capabilities and their cell-only resolution due to signal diffusion (Schulte et al., 2024; Wu et al., 2022). Additionally, lower entry costs and robustness of the canonical techniques makes them indispensable for initial research in obtaining the ground-truth pattern of gene expression.

Regardless of the ISH method, a new user will face the fact that every approach is tailored for specific antisense target-binding probes, which mainly fit only one of the methods. This thus requires either locking-in to a particular technique or substantial resource investments to accommodate several methods in one project. Additionally, when using commercial solutions, the exact probes’ design can sometimes be hidden from the end user, obscuring the research data and hindering troubleshooting and independent design of the study. Challenged by these issues, we looked for open, non-commercial alternatives that allow us to combine the benefits of canonical ISH approaches and modern semi-proprietary methods. We found that single-stranded DNA (ssDNA) probes with *in vitro* elongated concatemers produced by the signal amplification by exchange reaction (SABER) FISH protocol (Kishi et al., 2019) can be simply and universally applied for canonical colorimetric ISH and FISH methods as well as for the more recent enzyme-free one-step hybridization chain reaction (HCR) multiplex FISH (Choi et al., 2014; Choi et al., 2018; Schulte et al., 2024). Here, we demonstrate a “one probe fits all” approach, OneSABER, where a once-ordered set of generic user-defined short ssDNA oligonucleotides forms a platform for adaptable RNA (F)ISH signal amplification strength via SABER concatemers with customizable length. OneSABER is suitable for all commonly used detection methods.

The free-living marine flatworm *Macrostomum lignano* is a relatively new invertebrate model for studying regeneration, *in vivo* stem cell potency and differentiation, ageing, genome duplication and karyotype evolution, sexual selection, and many more biological questions (De Miguel-Bonet et al., 2018; Egger et al., 2006; Giannakara et al., 2016; Grudniewska et al.; Mouton et al., 2018; Patlar et al., 2020; Pfister et al., 2008; Santhosh et al., 2024; Ustyantsev and Berezikov, 2021; Ustyantsev et al., 2021a; Wudarski et al., 2020; Wunderer et al., 2019; Zadesenets et al., 2017; Zadesenets et al., 2023). Amenability to crucial molecular biology techniques such as RNA interference and, most importantly, transgenesis, combined with convenient laboratory culturing conditions and features such as a transparent body, make *M. lignano* a powerful and increasingly popular model organism (Hall et al., 2024; Mouton et al., 2024; Ustyantsev et al., 2021b; Wudarski et al., 2017; Wudarski et al., 2020). However, RNA ISH in *M. lignano* has been limited mostly to canonical AP-based colorimetric assays using long hapten-labeled antisense RNA probes (Grudniewska et al., 2018; Grudniewska et al.; Kuales et al., 2011; Pfister et al., 2007; Pfister et al., 2008; Wudarski et al., 2017; Wunderer et al., 2019; Zhou et al., 2015), hindering the experimental potential of this flatworm. As a proof-of-concept implementation of the OneSABER ISH platform, we show its utility for RNA ISH and multiplex TSA and HCR FISH in whole-mount samples of *M. lignano*. Additionally, we provide a detailed protocol for the “one probe fits all” SABER whole-mount ISH in *M. lignano* with liquid-exchange mini-columns (Irvine, 2007) to minimize sample losses and shorten the overall experimental time. Importantly, the method and the protocol should be easily adaptable to whole-mount preparation in other models.

## RESULTS AND DISCUSSION

### Overview of the OneSABER platform principles and its components

We developed the “one probe fits all” OneSABER platform as a modular approach stemming from classical ISH protocols and several recently developed methods. The aim is to provide a unified but still versatile and fully user-controlled ISH experimental design. The principal operation of OneSABER is depicted in Fig. 1.

**Fig. 1.**
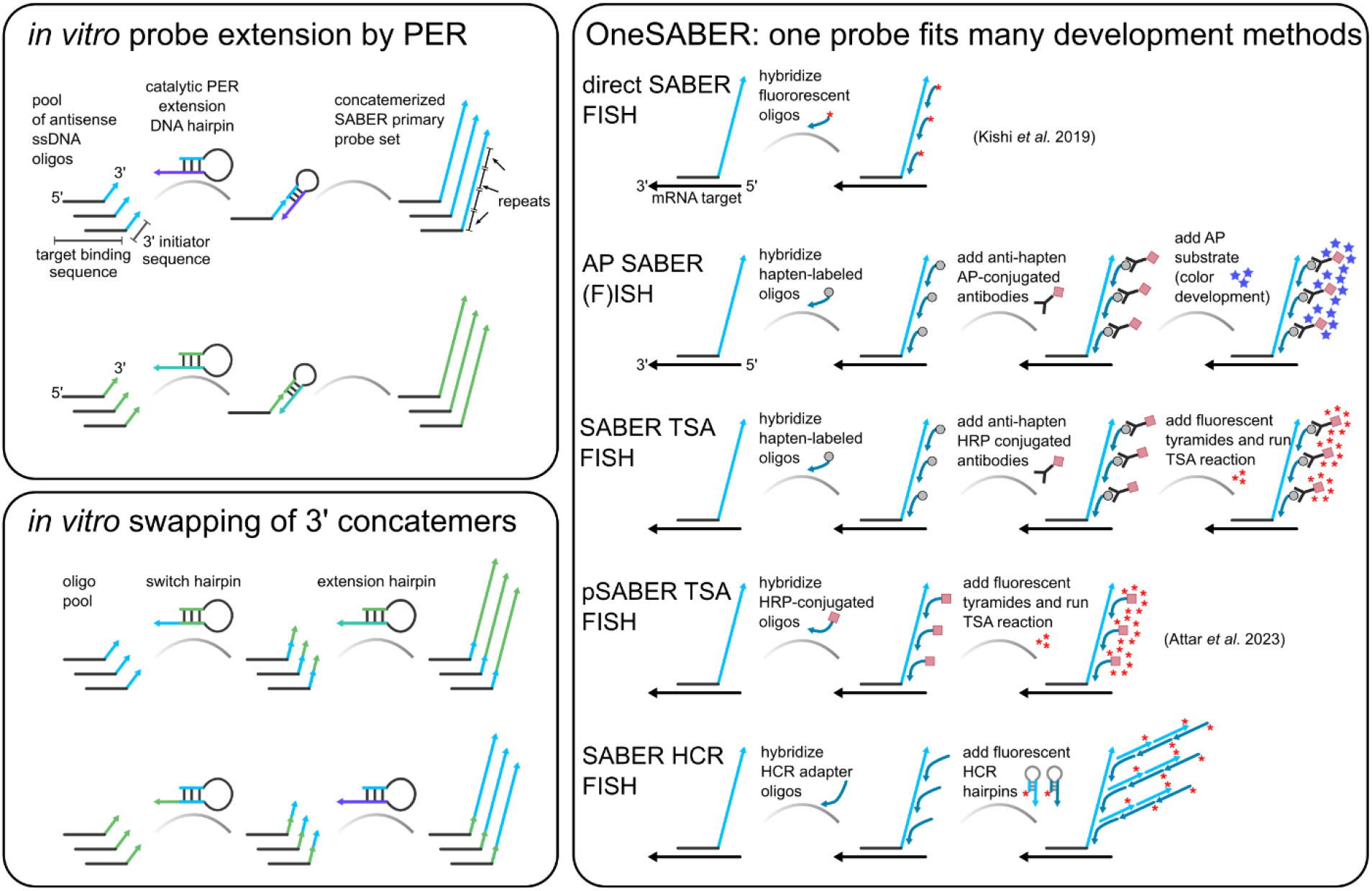
Principal operation scheme of the OneSABER mRNA *in situ* hybridization platform. Fluorophores are labeled as red asterisks. SABER - signal amplification by exchange reaction. PER – primer exchange reaction. AP – alkaline phosphatase. HRP – horse radish peroxidase. TSA – tyramide signal amplification. HCR – hybridization chain reaction. See Figs S1, S2 for additional details on probes/fluorophores switching and multiplexing.

In the heart of the platform is a pool of 15-30 custom user-defined short (~35-45 nt) ssDNA oligonucleotides complementary to an RNA target. The number of probes depends on the target RNA length and its expression strength (see Supplementary Information, Section S1). Each probe is ordered with a specific 9 nt 3’ initiator sequence and is extended *in vitro* through a primer exchange reaction (PER) (Kishi et al., 2018) (Supplementary Information, Section S1). PER relies on a catalytic DNA hairpin combined with a strand displacing polymerase and competitive branch migration to repeatedly add the same sequence to the 3’ end of the ssDNA primers, thus generating long concatemerized probes. The length of the extension is controlled by simply changing the reaction time and underlies the signal amplification strength (Kishi et al., 2019). The concatemers serve as universal landing pad sequences for binding short (20 nt) secondary oligonucleotide ssDNA probes/adapters, which are modified according to the chosen signal development method. Secondary probes labeled with hapten molecules like digoxigenin (DIG) and fluorescein (FITC/FAM) allow to apply standard enzyme-catalyzed reporter deposition assays, which are mediated by anti-hapten antibodies conjugated with AP or horse-radish peroxidase (HRP) for colorimetric and TSA fluorescent signal detection, respectively (Harper et al., 1997; King and Newmark, 2013; Paganos et al., 2022) Adding 5’ overhangs with HCR initiator sequences to the secondary probes converts them into adapters for fluorescent signal detection by HCR amplification (Choi et al., 2018; Wu et al., 2022) (Table S1). By changing the initiator sequence on the adapter probes, the same primary SABER probe pool can be targeted by different pairs of fluorescent HCR amplifiers, switching the detection channel (Fig. S1). Multiplexing is achieved by creating sets of primary SABER probes with different 3’ concatemer sequences, which, in turn, are specifically recognized by secondary probes with either different haptens (DIG, FAM, or dinitrophenol) or respective SABER-to-HCR adapters (Fig. S2, Table S1). Importantly, PER allows straightforward single-step switching of the concatemer sequences *in vitro* (Kishi et al., 2019), eliminating the need for ordering new sets of primary probe oligonucleotides (Fig. 1).

Apart from SABER FISH (Kishi et al., 2019), the OneSABER platform was inspired by several other studies. The idea of using haptenized secondary probes with subsequent antibody-mediated AP colorimetric substrate deposition came from a commercial BaseScope™ branched ISH/FISH protocol (Wang et al., 2019), while our SABER-to-HCR adapters are the result of rethinking of the π-FISH (Tao et al., 2023) and Yn-situ (Wu et al., 2022) adapter and pre-amplifier probes. Antibody-mediated SABER TSA is a modification of the peroxidase (p)SABER protocol published by the authors of SABER FISH (Attar et al., 2023), where they showed a substantial signal improvement when using directly HRP-conjugated secondary probes followed by TSA compared to the original protocol (Kishi et al., 2019).

### Colorimetric OneSABER *in situ* hybridization in *M. lignano*

In *M. lignano*, apart from a very recent HCR application (Hall et al., 2024), only results of traditional ISH/FISH assays using *in vitro* transcribed long antisense RNA probes with hapten-modified uridine bases have been reported (Grudniewska et al., 2018; Grudniewska et al.; Kuales et al., 2011; Lengerer et al., 2018; Mulder et al., 2010; Pfister et al., 2007; Pfister et al., 2008; Wudarski et al., 2017; Wunderer et al., 2019; Zhou et al., 2015). Compared to these RNA probes, short synthetic DNA probes provide several advantages, including user-controlled design of the most specific on-target sites, possibility to target short RNA transcripts (Tao et al., 2023), low hands-on processing time, and higher probe stability. We designed 7 sets of 15 to 31 ssDNA probes targeting genes expressed in the worm’s somatic tissues (neuronal *syt11*, intestinal *apob*, muscular *tnnt2*), testes (*boll* and *sperm1*), and proliferative cells including somatic stem cells and germline cells (*piwi and pcna*) (Table S2). The probe sets were concatemerized by the PER method to the approximate length of 170-300 nt (Table S3) and used for ISH experiments in *M. lignano* testing different signal development conditions. Our *M. lignano* ISH protocol contains several important modifications compared to the published traditional method (Pfister et al., 2007). From planarian and zebrafish protocols, we borrowed the combined fixation and permeabilization by adding acetic acid to the formaldehyde fixative, and we added a permeabilization step with hydrogen peroxide (Fernández and Fuentes, 2013; Gaetano and King, 2023; King and Newmark, 2013; Lauter et al., 2011). Hybridization buffers, wash solutions and hybridization temperatures were adapted to match the requirements of the short binding sequences of the primary and secondary SABER probes. Finally, we found that the use of liquid-exchange mini-columns (Irvine, 2007) instead of microcentrifuge tubes or multiwell plates during pre- and post-hybridization steps can prevent loss of smaller animals as well as make the exchange of hybridization and wash solutions faster, more convenient, and less wasteful (Supplementary Information, Section S2).

First, we tested colorimetric signal development with AP SABER on adult homeostatic (uncut) *M. lignano* worms using two different AP substrates: Vector Blue (Vector Labs) and nitro-blue tetrazolium chloride/5-bromo-4-chloro-3’-indolyphosphate p-toluidine (NBT/BCIP) (Fig. 2). Already after 10 min of development, strong and specific signals could be detected for the *boll, apob*, and *sperm1* genes, for *syt11, tnnt* and *piwi* it took around 40 min, and almost 2 hours for *pcna* (Fig. 2, Table S3). In some cases, however, we noticed recurring non-specific signal around the worms’ mouth and pharynx, reminiscent of the position of the *M. lignano* secretion cells in the head (Lengerer et al., 2016). This background is independent from the used SABER concatemer sequence and probe set. The pattern looks specific for the SABER protocol and differs from the standard ISH background in the gut and/or pharyngeal glands frequently observed with long antisense RNA probes in *M. lignano* (Lengerer et al., 2018; Pfister et al., 2007). In later experiments, we noticed that non-specific signal is completely absent when using younger worms (less than 6 weeks old) and shorter (< 500 nt) extended primary probes. *Syt11* (neuronal), *tnnt2* (muscle), and *pcna* (neoblasts) ISH patterns were obtained for the first time in *M. lignano*, while the patterns for the rest of the genes correspond to those previously published (Kuales et al., 2011; Pfister et al., 2007; Pfister et al., 2008; Wudarski et al., 2017). The *pcna* expression strongly resembles the pattern of *piwi* (Grudniewska et al.; Pfister et al., 2007), and the *tnnt2* pattern is similar to the previously published pattern of *myh6* (Wudarski et al., 2017). The pattern of *syt11* corresponds to the described anatomy of the *M. lignano* neural system (Hall et al., 2024; Ladurner et al., 1997; Ladurner et al., 2005; Morris et al., 2007).

**Fig. 2.**
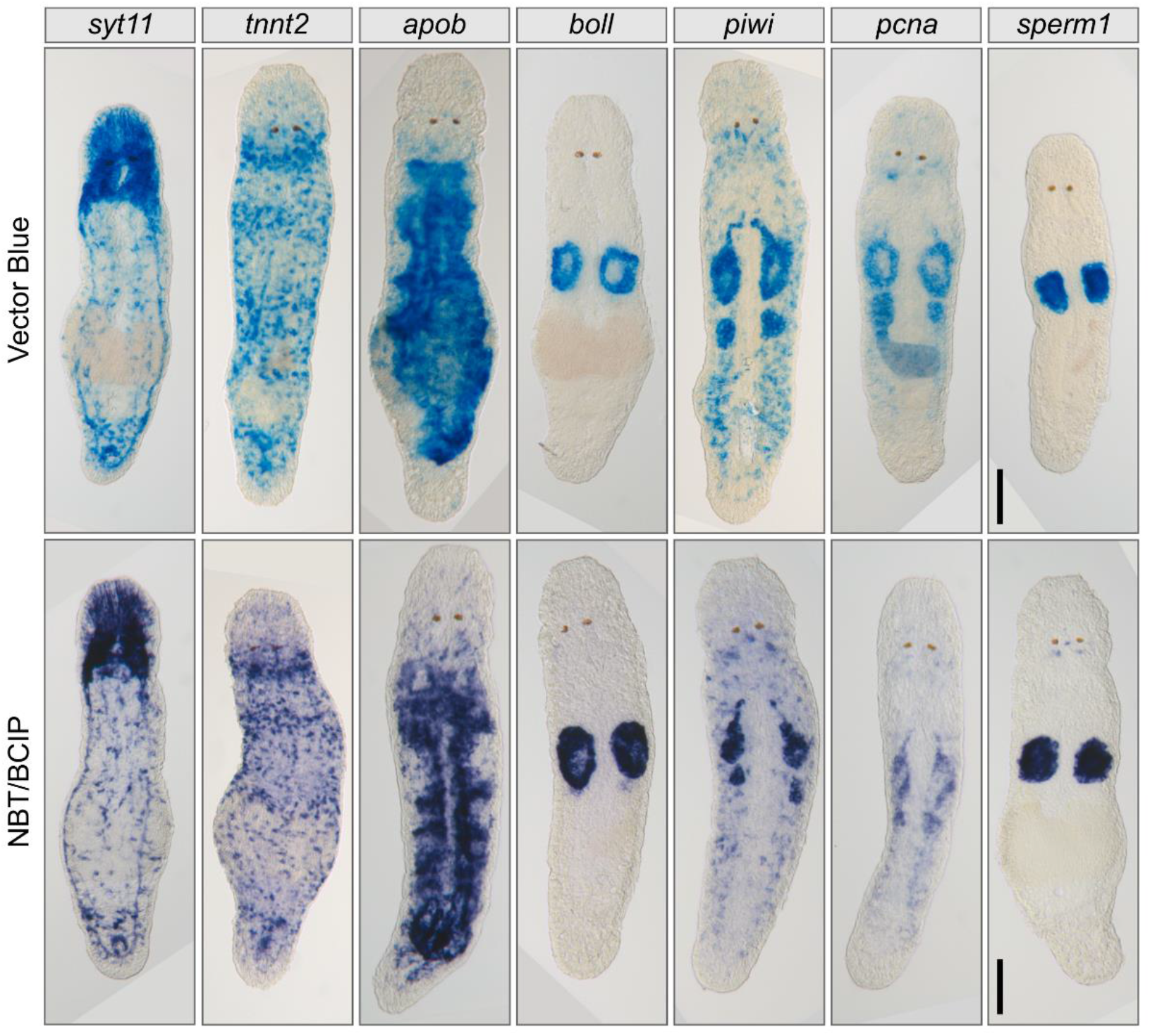
An example of alkaline phosphatase (AP) SABER colorimetric signal development on whole-mount *M. lignano* samples using different AP substrates. Vector Blue (Vector Labs) can also be viewed by fluorescent microscopy (Fig. S3). NBT/BCIP - nitro-blue tetrazolium chloride/5-bromo-4-chloro-3’-indolyphosphate p-toluidine salt. See Tables S2, S3 for development times and number of probes used for each mRNA target. The scale bars are 50 µm.

Colorimetric single gene ISHs should serve as ground truth for finding correct gene expression, estimating the sensitivity and specificity of the ordered probes, and should show if adjustments to the concatemer length are necessary before proceeding to less sensitive FISH assays. Samples can be stored for archiving indefinitely in any common mounting media. Vector Blue precipitates, like Fast Blue (Brown and Pearson, 2015; Lauter et al., 2011), can also be visualized fluorescently (Fig. S3) in the near far-red light spectrum and scanned by confocal microscopes without photobleaching by even strong laser powers. Additionally, these fluorescent substrates can be multiplexed with other FISH development assays such as TSA (Gaetano and King, 2023; Lauter et al., 2011; Schumacher et al., 2014).

### Fluorescent and multiplexed OneSABER *in situ* hybridization in *M. lignano*

On fixed cells and thin tissue sections, the original unbranched version of SABER FISH showed high flexibility and ease of multiplexing by using directly fluorophore-labeled secondary probes, called imagers (Kishi et al., 2019). This approach was validated by several independent studies (Amamoto et al., 2019; Chrysostomou et al.; Wang et al., 2020). However, when we tried to apply the original direct SABER FISH on whole-mount preparations of *M. lignano*, it resulted in low fluorescent intensity levels. This was even the case for strongly expressed targets like *syt11* with the highest number of ordered probes (Fig. S4, Table S2). For weaker targets, like *piwi*, the signal is barely detected even by confocal microscopy (Fig. S4). Lower signal in the absence of amplification through hybridization of secondary and tertiary branch probes was also acknowledged by the authors of SABER FISH (Attar et al., 2023). Branching would make the protocol longer and more complicated, while ordering more primary probes would be costly, and additional extension of concatemers may compromise their penetration into cells and increase background labeling. As a solution, Attar et al. developed the pSABER TSA protocol (Attar et al., 2023). We successfully tested this protocol, which uses secondary probes directly conjugated with HRP enzymes, making it faster than antibody-mediated SABER TSA (Fig. S5A). However, synthesis of HRP-conjugated oligonucleotides is not commonly offered and will be less accessible for most users. It also requires high initial investments with limited return (Tables S4, S5). Furthermore, in the OneSABER platform, it would be more beneficial to stay with antibody-mediated SABER TSA as it shares the same hapten-modified secondary probes to the AP SABER assay (Fig. 1). Importantly, fluorescent tyramides can be synthesized in-house and were shown to be efficient for TSA FISH in whole mount zebrafish and planarian samples (King and Newmark, 2013; Lauter et al., 2011), thus allowing significant potential cost reductions (Tables S4, S5) compared to commercial alternatives.

Therefore, we next tested SABER TSA FISH for the simultaneous detection of gene pairs of *syt11* with *apob* on adult uncut worms (Fig. 3A) and *piwi* and *boll* on regenerating worms (Fig. 3B) after tail plate amputation. Regeneration was employed to boost the *piwi* mRNA levels in somatic neoblasts and to highlight its expression in the blastema. TSA allows flexible fluorescent channel selection for each gene in a multiplex assay by simply changing fluorescent tyramides for development (Fig. 3, Fig. S5A). Overall, TSA SABER resulted in high signal-to-noise ratio FISH expression patterns with the whole cell cytoplasm evenly illuminated for both gene pairs.

**Fig. 3.**
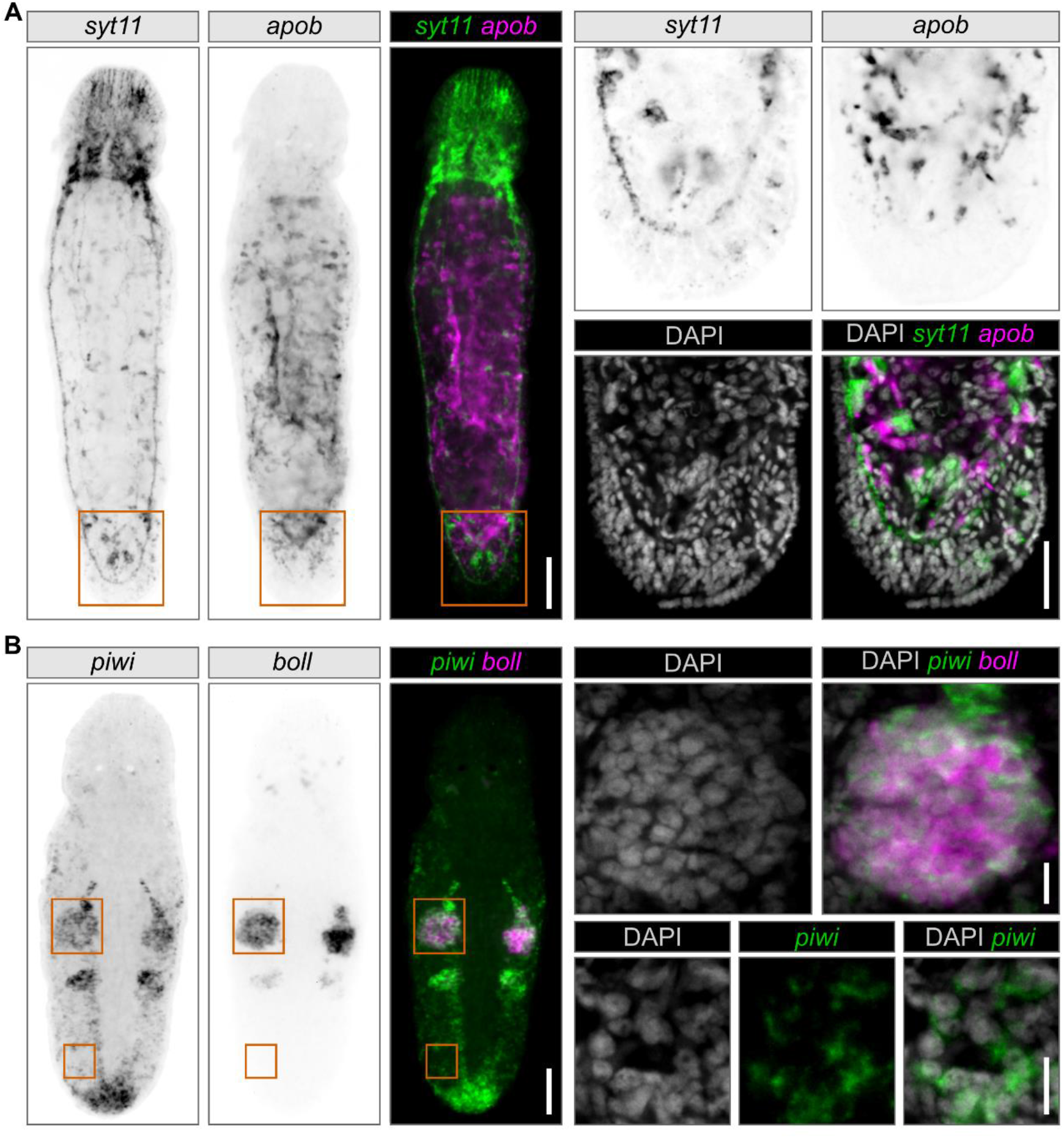
An example of duplex fluorescent signal development using SABER TSA on whole-mount homeostatic and regenerating *M. lignano* samples. The three panels on the left are widefield fluorescent microscopy photos. Grayscale panels are inverted images of different fluorescent channels provided for unbiased patterns’ comparison. Zoom-in regions are outlined in orange boxes and shown as panels on the right. Zoom-ins are average intensity z-projections of several confocal microscopy sections. (A) Expression patterns for a pair of somatic genes - *syt11* (neuronal) and *apob* (intestinal) in a homeostatic worm. Zoom-ins are for the tail plate region. *Apob* and s*yt11* were developed with fluorescein (FITC) and CF568 tyramides (Biotium), respectively. The scale bars for the left and right panels are 50 µm and 25 µm, respectively. (B) Expression patterns for a pair of germline and stem cell genes - *boll* (testes) and *piwi* (germline and stem cells) in a regenerating worm fixed 24 h post tail plate amputation. Zoom-ins are for the testis and somatic neoblasts. *Piwi* and *boll* were developed with FITC and CF647 tyramides (Biotium), respectively. The scale bars for the left and right panels are 50 µm and 10 µm, respectively.

Finally, we tested if SABER can be used in combination with HCR hairpin amplifiers. We combined probes for *syt11, tnnt*, and *apob* for SABER HCR on adult homeostatic worms (Fig. 4a) and *boll, piwi*, and *sperm1* on regenerating animals (Fig. 4b). The samples showed obvious tissue-specific fluorescent signals for each gene, although the patterns look finer outlined than in SABER TSA (Fig. 3) and appear like small grainy foci within the cells rather than evenly cytoplasm-diffused TSA fluorescence. As a result, despite higher intracellular resolution, some narrow anatomical structures like neural chords (*syt11*) or muscles (*tnnt2*) seem less visible for the human eye after SABER HCR (Figs. 3, 4). It is hard to compare brightness and strength of different fluorophores, especially in the context of different signal diffusion. When choosing between SABER HCR vs SABER TSA, their multiplexing potential must also be considered. Signal development in HCR happens simultaneously for all channels in a single step. In TSA, it is sequential and takes *N*x more time to develop for *N* different fluorescent signals than in HCR. Therefore, if TSA chemistry and antibodies are not yet available for a user and multiplexing is prioritized over signal strength, then investing in HCR chemistry over TSA might be a good solution (Tables S4, S5).

**Fig. 4.**
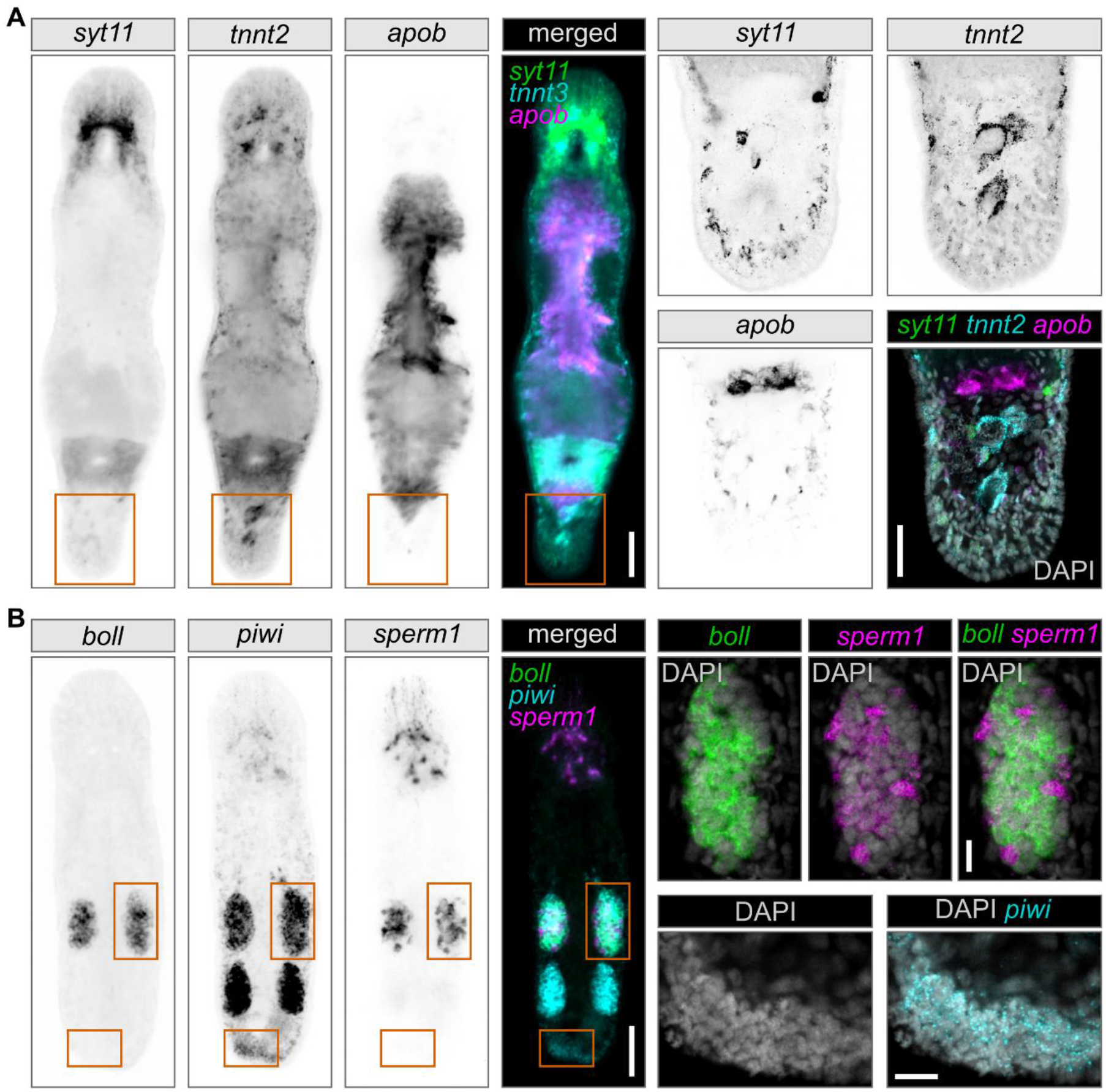
An example of triple multiplex fluorescent signal development using SABER HCR on whole-mount homeostatic and regenerating *M. lignano* samples. The four panels on the left are widefield fluorescent microscopy photos. Grayscale panels are inverted images of different fluorescent channels provided for unbiased patterns’ comparison. Fluorophores from left to right: Alexa488, Alexa546, and Alexa647. Zoom-in regions are outlined in orange boxes and shown on the right. Zoom-ins are average intensity z-projections of several confocal microscopy sections. (A) Expression patterns for three somatic genes - *syt11* (neuronal), *tnnt2* (muscular), and *apob* (intestinal) in a homeostatic worm. Zoom-ins are for the tail plate region. The scale bars for the left and right panels are 50 µm and 25 µm, respectively. (B) Expression patterns for three germline and stem cell genes - *boll* (testes), *piwi* (germline and stem cells), and *sperm1* (testes) in a regenerating worm fixed 24 h post tail plate amputation. Zoom-ins are for the testis and blastema regions. The scale bars for the left and right panels are 50 µm and 10 µm, respectively.

The current version of the HCR protocol relies on short split dual primary DNA probes, which were shown to significantly suppress the background HCR amplification from non-specific probe binding (Choi et al., 2018; Schulte et al., 2024). In principle, the primary split-probe approach is also compatible with the OneSABER platform, but we have not tested whether it will be beneficial in this setup. In SABER HCR, primary probes are approximately 10 times longer, may penetrate less into the sample, and cause unwanted background. However, apart from the already mentioned inherent non-specific binding to the areas in the head, SABER HCR showed a similar signal specificity to SABER TSA, indicating the viability of our single-probe approach for whole-mount samples of *M. lignano*.

There are two major advantages of SABER HCR over the original HCR protocol. First, similarly to the Yn-situ (Wu et al., 2022) and π-FISH (Tao et al., 2023) methods, longer landing-pad concatemer length of SABER probes should provide scalable linear signal pre-amplification and allow using fewer probes, thus saving costs. Second, one can seamlessly adapt the same SABER probe to any of the HCR hairpin amplifiers by simply changing the 5’ initiator sequence of the corresponding SABER-to-HCR adapters. This eliminates the need to order multiple sets of HCR amplifiers labeled by different fluorophores, since any SABER probe can be used with any of the HCR hairpins. Ordering only a single set of HCR hairpins substantially reduces costs for planning complex experiments with fluorescent channel switching. To demonstrate this, we fully swapped the fluorescence channels for the same probes of *sperm1, syt11*, and *piwi* in another SABER HCR experiment, where we also re-used a mix of annealed HCR amplifiers from the previous experiments (Fig. S5B).

In conclusion, we demonstrated the utility of the “one probe fits all” OneSABER platform on whole mount preparations of homeostatic and regenerating *M. lignano* flatworms. For this, we also developed and validated a new fixation, permeabilization, and hybridization protocol for *M. lignano* adapted for the use of short target binding sequences of SABER probes with long concatemers, faster and loss-resistant sample handling by liquid-exchange mini columns. We summarized the advantages and most suitable applications for each of the signal development protocols of OneSABER in Table S6. The ISH protocol and the platform operation principles shown here should be easily transferable to other animal research models and equip the researchers with a robust toolkit for fine-tuned ISH experiments.

## MATERIALS AND METHODS

### M. lignano culturing and sample preparation for ISH

*M. lignano* wild-type NL12 worms were cultured at standard conditions at 20°C as previously described (Mouton et al., 2024). For the ISH experiments, 3–5-week-old sexually mature animals were selected and prepared for fixation according to the protocol detailed in Supplementary Information, Section S4.1. For regeneration condition, the tail of worms of the same age was amputated below the ovaries, worms were left to regenerate for 24 hours without food, and were then fixed as in Supplementary Information, Section S4.1.

### Probes, PER concatemerization and HCR amplifiers

Except for 5’ peroxidase-conjugated secondary probes for TSA SABER FISH (Eurogentec) and HCR amplifiers for SABER-HCR FISH (Molecular Instruments), all DNA oligonucleotides were ordered from Integrated DNA technologies. All probe sequences were ordered in a 25 nmol 96-well standard desalting format to save costs. Sequences, applications, and purification methods for each oligonucleotide are specified in Tables S1 and S2. Antisense ssDNA probes with 3 different 3’ PER initiator sequences (p27, p28, and p30) (Kishi et al., 2019) were designed, checked *in silico* for target specificity, *in vitro* concatemerized to 200-500 nt by PER, and purified as detailed in Supplementary Information, Section S1.

### In situ hybridization and signal detection

A detailed step-by-step protocol, including buffer recipes, used antibodies, fluorescent tyramides, as well as mounting of the animals for microscopy is provided in Supplementary Information, Sections S3-S4. Samples developed with colorimetric AP SABER were mounted in 80% glycerol/PBStw, while FISH samples were mounted in VECTASHIELD Vibrance (Vector Labs).

### Microscopy

Widefield microscopy photos were made using a Zeiss Axio Zoom V.16 stereo fluorescent microscope equipped with a PlanNeoFluar Z 2.3x/0.57 objective, HXP 200W lamp, and Zeiss filter sets 96HE (“blue”, BP390/40, BP450/40), 38HE (“green”, BP470/40, BP525/50), 43HE (“red”, BP550/25, BP605/70), and filter set 50 (“far-red”, BP640/30, BP690/50). Colorimetric ISH photos were taken with an AxioCam MRc5 color camera, and fluorescent images with an AxioCam HRm CCD camera. Confocal microscopy photos were obtained with a Leica SP8x equipped with a HC PL APO CS2 63x/1.4 oil objective at the UMCG Imaging and Microscopy Center (UMIC). Processing of raw microscopy images was done in Fiji ImageJ (Schindelin et al., 2012). Final figure panels were assembled in Inkscape v. 1.3.2 (https://inkscape.org).

## ACKNOWLEDGEMENTS

We thank Dr. Erin Davies (NIH) for sharing her lab’s unpublished HCR data and reagents and providing an online seminar platform for fruitful discussions between the *Macrostomum* research groups from the US and the Netherlands, thus inspiring us to try combining HCR with SABER.

## AUTHOR CONTRIBUTIONS

Conceptualization: KU, SM, EB; Methodology: KU, FJV; Validation: KU, FJV, MS; Investigation: KU, FJV, MS; Writing - original draft: KU, MS, SM, EB; Writing - review & editing: KU, MS, SM, EB; Visualization: KU; Supervision: EB; Funding acquisition: EB.

## REFERENCES

Attar, S., Browning, V. E., Liu, Y., Nichols, E. K., Tsue, A. F., Shechner, D. M., Shendure, J., Lieberman, J. A., Akilesh, S. and Beliveau, B. J. (2023). Efficient and highly amplified imaging of nucleic acid targets in cellular and histopathological samples with pSABER. 2023.01.30.526264.

Bobrow, M. N., Harris, T. D., Shaughnessy, K. J. and Litt, G. J. (1989). Catalyzed reporter deposition, a novel method of signal amplification. Application to immunoassays. J. Immunol. Methods 125, 279–285.

Brown, D. D. R. and Pearson, B. J. (2015). One FISH, dFISH, Three FISH: Sensitive Methods of Whole-Mount Fluorescent In Situ Hybridization in Freshwater Planarians. In In Situ Hybridization Methods (ed. Hauptmann, G.), pp. 127–150. New York, NY: Springer.

Chari, T., Weissbourd, B., Gehring, J., Ferraioli, A., Leclère, L., Herl, M., Gao, F., Chevalier, S., Copley, R. R., Houliston, E., et al. (2021). Whole-animal multiplexed single-cell RNA-seq reveals transcriptional shifts across Clytia medusa cell types. Sci. Adv. 7, eabh1683.

Choi, H. M. T., Beck, V. A. and Pierce, N. A. (2014). Next-Generation in Situ Hybridization Chain Reaction: Higher Gain, Lower Cost, Greater Durability. ACS Nano 8, 4284–4294.

Choi, H. M. T., Schwarzkopf, M., Fornace, M. E., Acharya, A., Artavanis, G., Stegmaier, J., Cunha, A. and Pierce, N. A. (2018). Third-generation in situ hybridization chain reaction: multiplexed, quantitative, sensitive, versatile, robust. Dev. Camb. Engl. 145, dev165753.

Dardani, I., Emert, B. L., Goyal, Y., Jiang, C. L., Kaur, A., Lee, J., Rouhanifard, S. H., Alicea, G. M., Fane, M. E., Xiao, M., et al. (2022). ClampFISH 2.0 enables rapid, scalable amplified RNA detection in situ. Nat. Methods 19, 1403–1410.

De Miguel-Bonet, M. D. M., Ahad, S. and Hartenstein, V. (2018). Role of neoblasts in the patterned postembryonic growth of the platyhelminth Macrostomum lignano. Neurogenesis 5, e1469944-1-e1469944-9.

Egger, B., Ladurner, P., Nimeth, K., Gschwentner, R. and Rieger, R. (2006). The regeneration capacity of the flatworm Macrostomum lignano--on repeated regeneration, rejuvenation, and the minimal size needed for regeneration. Dev. Genes Evol. 216, 565–577.

Fernández, J. and Fuentes, R. (2013). Fixation/permeabilization: new alternative procedure for immunofluorescence and mRNA in situ hybridization of vertebrate and invertebrate embryos. Dev. Dyn. Off. Publ. Am. Assoc. Anat. 242, 503–517.

Fincher, C. T., Wurtzel, O., de Hoog, T., Kravarik, K. M. and Reddien, P. W. (2018). Cell type transcriptome atlas for the planarian Schmidtea mediterranea. Science 360, eaaq1736.

Gaetano, A. J. and King, R. S. (2023). A simplified and rapid in situ hybridization protocol for planarians. BioTechniques 75, 231–239.

Giannakara, A., Schärer, L. and Ramm, S. A. (2016). Sperm competition-induced plasticity in the speed of spermatogenesis. BMC Evol. Biol. 16, 60.

Goh, J. J. L., Chou, N., Seow, W. Y., Ha, N., Cheng, C. P. P., Chang, Y.-C., Zhao, Z. W. and Chen, K. H. (2020). Highly specific multiplexed RNA imaging in tissues with split-FISH. Nat. Methods 17, 689– 693.

Grudniewska, M., Mouton, S., Grelling, M., Wolters, A. H. G., Kuipers, J., Giepmans, B. N. G. and Berezikov, E. (2018). A novel flatworm-specific gene implicated in reproduction in Macrostomum lignano. Sci. Rep. 8, 3192.

Grudniewska, M., Mouton, S., Simanov, D., Beltman, F., Grelling, M., de Mulder, K., Arindrarto, W., Weissert, P. M., van der Elst, S. and Berezikov, E. Transcriptional signatures of somatic neoblasts and germline cells in Macrostomum lignano. eLife 5, e20607.

Hadwiger, G., Dour, S., Arur, S., Fox, P. and Nonet, M. L. (2010). A monoclonal antibody toolkit for C. elegans. PloS One 5, e10161.

Hall, N., Li, H., Chai, C., Vermeulen, S., Bigasin, R., Song, E. S., Gibson, J., Prakash, M., Fire, A. and Wang, B. (2024). A genetic and microscopy toolkit for live imaging of whole-body regeneration in<i>Macrostomum lignano</i.

Harper, S. J., Bailey, E., McKeen, C. M., Stewart, A. S., Pringle, J. H., Feehally, J. and Brown, T. (1997). A comparative study of digoxigenin, 2, 4-dinitrophenyl, and alkaline phosphatase as deoxyoligonucleotide labels in non-radioisotopic in situ hybridisation. J. Clin. Pathol. 50, 686– 690.

Hemmati-Brivanlou, A., Frank, D., Bolce, M. E., Brown, B. D., Sive, H. L. and Harland, R. M. (1990). Localization of specific mRNAs in Xenopus embryos by whole-mount in situ hybridization. Dev. Camb. Engl. 110, 325–330.

Hulett, R. E., Kimura, J. O., Bolaños, D. M., Luo, Y.-J., Rivera-López, C., Ricci, L. and Srivastava, M. (2023). Acoel single-cell atlas reveals expression dynamics and heterogeneity of adult pluripotent stem cells. Nat. Commun. 14, 2612.

Irvine, S. Q. (2007). Whole-mount in situ hybridization of small invertebrate embryos using laboratory mini-columns. BioTechniques 43, 764–768.

Jowett, T. and Lettice, L. (1994). Whole-mount in situ hybridizations on zebrafish embryos using a mixture of digoxigenin- and fluorescein-labelled probes. Trends Genet. TIG 10, 73–74.

King, R. S. and Newmark, P. A. (2013). In situ hybridization protocol for enhanced detection of gene expression in the planarian Schmidtea mediterranea. BMC Dev. Biol. 13, 8.

Kishi, J. Y., Schaus, T. E., Gopalkrishnan, N., Xuan, F. and Yin, P. (2018). Programmable autonomous synthesis of single-stranded DNA. Nat. Chem. 10, 155–164.

Kishi, J. Y., Lapan, S. W., Beliveau, B. J., West, E. R., Zhu, A., Sasaki, H. M., Saka, S. K., Wang, Y., Cepko, C. L. and Yin, P. (2019). SABER amplifies FISH: enhanced multiplexed imaging of RNA and DNA in cells and tissues. Nat. Methods 16, 533–544.

Kuales, G., De Mulder, K., Glashauser, J., Salvenmoser, W., Takashima, S., Hartenstein, V., Berezikov, E., Salzburger, W. and Ladurner, P. (2011). Boule-like genes regulate male and female gametogenesis in the flatworm Macrostomum lignano. Dev. Biol. 357, 117–132.

Ladurner, P., Mair, G. R., Reiter, D., Salvenmoser, W. and Rieger, R. M. (1997). Serotonergic Nervous System of Two Macrostomid Species: Recent or Ancient Divergence? Invertebr. Biol. 116, 178– 191.

Ladurner, P., Pfister, D., Seifarth, C., Schärer, L., Mahlknecht, M., Salvenmoser, W., Gerth, R., Marx, F. and Rieger, R. (2005). Production and characterisation of cell- and tissue-specific monoclonal antibodies for the flatworm Macrostomum sp. Histochem. Cell Biol. 123, 89–104.

Lauter, G., Söll, I. and Hauptmann, G. (2011). Two-color fluorescent in situ hybridization in the embryonic zebrafish brain using differential detection systems. BMC Dev. Biol. 11, 43.

Lengerer, B., Hennebert, E., Flammang, P., Salvenmoser, W. and Ladurner, P. (2016). Adhesive organ regeneration in Macrostomum lignano. BMC Dev. Biol. 16, 20.

Lengerer, B., Wunderer, J., Pjeta, R., Carta, G., Kao, D., Aboobaker, A., Beisel, C., Berezikov, E., Salvenmoser, W. and Ladurner, P. (2018). Organ specific gene expression in the regenerating tail of Macrostomum lignano. Dev. Biol. 433, 448–460.

Luehrsen, K. R., Davidson, S., Lee, Y. J., Rouhani, R., Soleimani, A., Raich, T., Cain, C. A., Collarini, E. J., Yamanishi, D. T., Pearson, J., et al. (2000). High-density hapten labeling and HRP conjugation of oligonucleotides for use as in situ hybridization probes to detect mRNA targets in cells and tissues. J. Histochem. Cytochem. Off. J. Histochem. Soc. 48, 133–145.

Morris, J., Cardona, A., De Miguel-Bonet, M. D. M. and Hartenstein, V. (2007). Neurobiology of the basal platyhelminth Macrostomum lignano: map and digital 3D model of the juvenile brain neuropile. Dev. Genes Evol. 217, 569–584.

Mouton, S., Grudniewska, M., Glazenburg, L., Guryev, V. and Berezikov, E. (2018). Resilience to aging in the regeneration-capable flatworm Macrostomum lignano. Aging Cell 17, e12739.

Mouton, S., Mougel, A., Ustyantsev, K., Dissous, C., Melnyk, O., Berezikov, E. and Vicogne, J. (2024). Optimized protocols for RNA interference in Macrostomum lignano. G3 GenesGenomesGenetics 14, jkae037.

Mulder, K. D., Kuales, G., Pfister, D., Egger, B., Seppi, T., Eichberger, P., Borgonie, G. and Ladurner, P. (2010). Potential of Macrostomum lignano to recover from γ-ray irradiation. Cell Tissue Res. 339, 527.

Paganos, P., Caccavale, F., La Vecchia, C., D’Aniello, E., D’Aniello, S. and Arnone, M. I. (2022). FISH for All: A Fast and Efficient Fluorescent In situ Hybridization (FISH) Protocol for Marine Embryos and Larvae. Front. Physiol. 13, 878062.

Patlar, B., Weber, M., Temizyürek, T. and Ramm, S. A. (2020). Seminal Fluid-Mediated Manipulation of Post-mating Behavior in a Simultaneous Hermaphrodite. Curr. Biol. CB 30, 143–149.e4.

Peter, A. K., Crocini, C. and Leinwand, L. A. (2017). Expanding our scientific horizons: utilization of unique model organisms in biological research. EMBO J. 36, 2311–2314.

Pfister, D., De Mulder, K., Philipp, I., Kuales, G., Hrouda, M., Eichberger, P., Borgonie, G., Hartenstein, V. and Ladurner, P. (2007). The exceptional stem cell system of Macrostomum lignano: Screening for gene expression and studying cell proliferation by hydroxyurea treatment and irradiation. Front. Zool. 4, 9.

Pfister, D., De Mulder, K., Hartenstein, V., Kuales, G., Borgonie, G., Marx, F., Morris, J. and Ladurner, P. (2008). Flatworm stem cells and the germ line: developmental and evolutionary implications of macvasa expression in Macrostomum lignano. Dev. Biol. 319, 146–159.

Santhosh, S., Ebert, D. and Janicke, T. (2024). Sperm competition favours intermediate sperm size in a hermaphrodite. J. Evol. Biol. voae058.

Schindelin, J., Arganda-Carreras, I., Frise, E., Kaynig, V., Longair, M., Pietzsch, T., Preibisch, S., Rueden, C., Saalfeld, S., Schmid, B., et al. (2012). Fiji: an open-source platform for biological-image analysis. Nat. Methods 9, 676–682.

Schmidt, B. F., Chao, J., Zhu, Z., DeBiasio, R. L. and Fisher, G. (1997). Signal amplification in the detection of single-copy DNA and RNA by enzyme-catalyzed deposition (CARD) of the novel fluorescent reporter substrate Cy3.29-tyramide. J. Histochem. Cytochem. Off. J. Histochem. Soc. 45, 365– 373.

Schulte, S. J., Fornace, M. E., Hall, J. K., Shin, G. J. and Pierce, N. A. (2024). HCR spectral imaging: 10-plex, quantitative, high-resolution RNA and protein imaging in highly autofluorescent samples. Dev. Camb. Engl. 151, dev202307.

Schumacher, J. A., Zhao, E. J., Kofron, M. J. and Sumanas, S. (2014). Two-color fluorescent in situ hybridization using chromogenic substrates in zebrafish. BioTechniques 57, 254–256.

Tao, Y., Zhou, X., Sun, L., Lin, D., Cai, H., Chen, X., Zhou, W., Yang, B., Hu, Z., Yu, J., et al. (2023). Highly efficient and robust π-FISH rainbow for multiplexed in situ detection of diverse biomolecules. Nat. Commun. 14, 443.

Tautz, D. and Pfeifle, C. (1989). A non-radioactive in situ hybridization method for the localization of specific RNAs in Drosophila embryos reveals translational control of the segmentation gene hunchback. Chromosoma 98, 81–85.

Toledano, H., D’Alterio, C., Loza-Coll, M. and Jones, D. L. (2012). Dual fluorescence detection of protein and RNA in Drosophila tissues. Nat. Protoc. 7, 1808–1817.

Ustyantsev, K. V. and Berezikov, E. V. (2021). Computational analysis of spliced leader trans-splicing in the regenerative flatworm Macrostomum lignano reveals its prevalence in conserved and stem cell related genes. Vavilovskii Zhurnal Genet. Sel. 25, 101–107.

Ustyantsev, K. V., Vavilova, V. Yu., Blinov, A. G. and Berezikov, E. V. (2021a). Macrostomum lignano as a model to study the genetics and genomics of parasitic flatworms. Vavilov J. Genet. Breed. 25, 108–116.

Ustyantsev, K., Wudarski, J., Sukhikh, I., Reinoite, F., Mouton, S. and Berezikov, E. (2021b). Proof of principle for piggyBac-mediated transgenesis in the flatworm Macrostomum lignano. Genetics 218, iyab076.

Wang, Y.-S. and Guo, J. (2021). Multiplexed Single-Cell in situ RNA Profiling. Front. Mol. Biosci. 8, 775410.

Wang, F., Flanagan, J., Su, N., Wang, L.-C., Bui, S., Nielson, A., Wu, X., Vo, H.-T., Ma, X.-J. and Luo, Y. (2012). RNAscope: a novel in situ RNA analysis platform for formalin-fixed, paraffin-embedded tissues. J. Mol. Diagn. JMD 14, 22–29.

Wang, Y., Xu, W., Maddera, L., Tsuchiya, D., Thomas, N., Yu, C. R. and Parmely, T. (2019). Alkaline phosphatase-based chromogenic and fluorescence detection method for BaseScopeTM In Situ hybridization. J. Histotechnol. 42, 193–201.

Wu, Y., Xu, W., Ma, L., Yu, Z., Wang, Y. and Yu, C. R. (2022). Robust and sensitive in situ RNA detection using Yn-situ. Cell Rep. Methods 2, 100201.

Wudarski, J., Simanov, D., Ustyantsev, K., de Mulder, K., Grelling, M., Grudniewska, M., Beltman, F., Glazenburg, L., Demircan, T., Wunderer, J., et al. (2017). Efficient transgenesis and annotated genome sequence of the regenerative flatworm model Macrostomum lignano. Nat. Commun. 8, 2120.

Wudarski, J., Egger, B., Ramm, S. A., Schärer, L., Ladurner, P., Zadesenets, K. S., Rubtsov, N. B., Mouton, S. and Berezikov, E. (2020). The free-living flatworm Macrostomum lignano. EvoDevo 11, 5.

Wunderer, J., Lengerer, B., Pjeta, R., Bertemes, P., Kremser, L., Lindner, H., Ederth, T., Hess, M. W., Stock, D., Salvenmoser, W., et al. (2019). A mechanism for temporary bioadhesion. Proc. Natl. Acad. Sci. 116, 4297–4306.

Xia, C., Babcock, H. P., Moffitt, J. R. and Zhuang, X. (2019). Multiplexed detection of RNA using MERFISH and branched DNA amplification. Sci. Rep. 9, 7721.

Young, A. P., Jackson, D. J. and Wyeth, R. C. (2020). A technical review and guide to RNA fluorescence in situ hybridization. PeerJ 8, e8806.

Zadesenets, K. S., Schärer, L. and Rubtsov, N. B. (2017). New insights into the karyotype evolution of the free-living flatworm Macrostomum lignano (Platyhelminthes, Turbellaria). Sci. Rep. 7, 6066.

Zadesenets, K. S., Ershov, N. I., Bondar, N. P. and Rubtsov, N. B. (2023). Unraveling the Unusual Subgenomic Organization in the Neopolyploid Free-Living Flatworm Macrostomum lignano. Mol. Biol. Evol. 40, msad250.

Zhou, X., Battistoni, G., El Demerdash, O., Gurtowski, J., Wunderer, J., Falciatori, I., Ladurner, P., Schatz, M. C., Hannon, G. J. and Wasik, K. A. (2015). Dual functions of Macpiwi1 in transposon silencing and stem cell maintenance in the flatworm Macrostomum lignano. RNA N. Y. N 21, 1885–1897.

