## Supplementary Information for "One probe fits all: a highly customizable modular RNA *in situ* hybridization platform expanding the application of SABER DNA probes"

### Contents

Tables S1–S6.......................................................................................pp. 2-11

Figs. S1–S7.........................................................................................pp. 12-16

Supplementary Materials and Methods (S1-S4):

*S1. Design, preparation, and validation of SABER probes*................p. 17

*S2. Assembly and operation of hybridization columns*....................p. 24

*S3. Recipes for the in situ hybridization protocol in M. lignano*.........p. 25

*S4. In situ hybridization protocol in M. lignano*...............................p. 28

References..........................................................................................p. 38

**Table S1.** Primer exchange reaction oligonucleotides, secondary hybridization probes, and HCR hairpins used in the study

| Oligo name | Oligo sequence (5' -> 3'), ordered from IDT unless specified otherwise | Purification |
|---|---|---|
| *Main hairpins for Primer Exchange Reaction (PER)* | | Standard desalting |
| Hairpin.p27_to_p27 | ACATCATCATGGGCCTTTTGGCCCATGATGATGTATGATGATGttttttt | |
| Hairpin.p28_to_p28 | ACAACTTAACGGGCCTTTTGGCCCGTTAAGTTGTGTTAAGTTGttttttt | |
| Hairpin.p30_to_p30 | AAATACTCTCGGGCCTTTTGGCCCGAGAGTATTTGAGAGTATTttttttt | |
| *Switching hairpins for Primer Exchange Reaction (PER)* | | |
| SwitchHairpin.p27_to_p28 | ACAACTTAACGGGCCTTTTGGCCCGTTAAGTTGTATGATGATGttttttt | |
| SwitchHairpin.p27_to_p30 | AAATACTCTCGGGCCTTTTGGCCC GAGAGTATTTATGATGATGttttttt | |
| SwitchHairpin.p28_to_p27 | ACATCATCATGGGCCTTTTGGCCCATGATGATGTGTTAAGTTGttttttt | |
| SwitchHairpin.p28_to_p30 | AAATACTCTCGGGCCTTTTGGCCCGAGAGTATTTGTTAAGTTGttttttt | |
| SwitchHairpin.p30_to_p27 | ACATCATCATGGGCCTTTTGGCCCATGATGATGTGAGAGTATTttttttt | |
| SwitchHairpin.p30_to_p28 | ACAACTTAACGGGCCTTTTGGCCCGTTAAGTTGTGAGAGTATTttttttt | |
| *Accesory oligonucliotide to absorbe potentially contaminating dGTPs from the in vitro reaction* | | |
| Clean.G | CCCCGAAAGTGGCCTCGGGCCTTTTGGCCCGAGGCCACTTTCG | |
| *Fluorophore-labeled secondary probes for direct SABER FISH* | | HPLC |
| p27-SABER-ATTO633 | /5ATTO633N/tt ATGATGATGT ATGATGATGT | |
| p30-SABER-ATTO488 | /5ATTO488N/tt GAGAGTATT T GAGAGTATT T | |
| *Hapten-conjugated secondary probes for AP SABER and SABER TSA* | | HPLC |
| p27-SABER-DIG | ATGATGATGtATGATGATGt tttttttt/3DIG_N/ | |
| p28-SABER-6FAM | GTTAAGTTGtGTTAAGTTGt tttttttt/36-FAM/ | Standard Desalting |
| p30-SABER-6FAM | GAGAGTATTtGAGAGTATTt tttttttt/36-FAM/ | |
| *SABER to HCR adapters* (concatemers to initiators*) | | Standard Desalting |
| p27-SABER_to_B1-HCR | GAGGAGGGCAGCAAACGGGAAGAGTCTTCCTTTACGttttATGATGATGtATGATGATGt | |
| p28-SABER_to_B1-HCR | GAGGAGGGCAGCAAACGGGAAGAGTCTTCCTTTACGttttGTTAAGTTGtGTTAAGTTGt | |
| p30-SABER_to_B1-HCR | GAGGAGGGCAGCAAACGGGAAGAGTCTTCCTTTACGttttGAGAGTATTtGAGAGTATTt | |
| p27-SABER_to_B2-HCR | CCTCGTAAATCCTCATCAATCATCCAGTAAACCGCCttttATGATGATGtATGATGATGt | |
| p28-SABER_to_B2-HCR | CCTCGTAAATCCTCATCAATCATCCAGTAAACCGCCttttGTTAAGTTGtGTTAAGTTGt | |
| p30-SABER_to_B2-HCR | CCTCGTAAATCCTCATCAATCATCCAGTAAACCGCCttttGAGAGTATTtGAGAGTATTt | |
| p27-SABER_to_B3-HCR | GTCCCTGCCTCTATATCTCCACTCAACTTTAACCCGttttATGATGATGtATGATGATGt | |
| p28-SABER_to_B3-HCR | GTCCCTGCCTCTATATCTCCACTCAACTTTAACCCGttttGTTAAGTTGtGTTAAGTTGt | |
| p30-SABER_to_B3-HCR | GTCCCTGCCTCTATATCTCCACTCAACTTTAACCCGttttGAGAGTATTtGAGAGTATTt | |
| *Directly peroxidase-conjugated secondaty probes for pSABER TSA (Ordered from Eurogentec)* |  | IEX-FPLC |
| p27-SABER-5'HRP | [HRP Conjugation]tttttttttATGATGATGTATGATGATG | |
| p28-SABER-5'HRP | [HRP Conjugation]tttttttttGTTAAGTTGTGTTAAGTTG | |
| *HCR hairpin amplifiers (ordered from Molecular Instruments)* | | |
| B1-H1 | CGTAAAGGAAGACTCTTCCCGTTTGCTGCCCTCCTCGCATTCTTTCTTGAGGAGGGCAGCAAACGGGAAGAG /C9-Alexa Fluor 488-3’/ | |
| B1-H2 | /5’-Alexa Fluor 488-C12/ GAGGAGGGCAGCAAACGGGAAGAGTCTTCCTTTACGCTCTTCCCGTTTGCTGCCCTCCTCAAGAAAGAATGC | |

| | |
|---|---|
| B2-H1 | GGCGGTTTACTGGATGATTGATGAGGATTTACGAGGAGCTCAGTCCATCCTC GTAAATCCTCATCAATCATC /C9-Alexa Fluor 546-3’/ |
| B2-H2 | /5’-Alexa Fluor 546-C12/ CCTCGTAAATCCTCATCAATCATCCAGTAAACCGCCGATGATTGATGAGGAT TTACGAGGATGGACTGAGCT |
| B3-H1 | CGGGTTAAAGTTGAGTGGAGATATAGAGGCAGGGACAAAGTCTAATCCGTC CCTGCCTCTATATCTCCACTC /C9-Alexa Fluor 647-3’/ |
| B3-H2 | /5’-Alexa Fluor 647-C12/ GTCCCTGCCTCTATATCTCCACTCAACTTTAACCCGGAGTGGAGATATAGAG GCAGGGACGGATTAGACTTT |

** HCR initiator sequences are underlined*

**Table S2.** Target genes with corresponding primary probe sequences used in the study

| ID | Probe target binding sequence (5' -> 3') | Ordered probe sequence (5' -> 3', from IDT) |
|---|---|---|
| | ***M. lignano* GeneID \| Gene Annotation \| Number of paralogs in the genome assembly \| PER 3' extension initiator sequence (5' -> 3')** | |
| | ***Mlig455_055907 \| boll \| 3 \| p27 (tttCATCATCAT)*** | |
| 1 | AAAATATTAACCCCAGCGTGCGCTCAACTCAACTCGCGCT | AAAATATTAACCCCAGCGTGCGCTCAACTCAACTCGCGCTtttCATCATCAT |
| 2 | AAAATCCTCGCGCGGATTGGCATCATTATTACACAGTTCTG | AAAATCCTCGCGCGGATTGGCATCATTATTACACAGTTCTGtttCATCATCAT |
| 3 | TAGCTAATTCCAGGCAATGCGTAAGGGAGAAGAACTTGGC | TAGCTAATTCCAGGCAATGCGTAAGGGAGAAGAACTTGGCtttCATCATCAT |
| 4 | AAAACAGTCGAATAGGACTGACGAAACAAGCATTTTGCTGCT | AAAACAGTCGAATAGGACTGACGAAACAAGCATTTTGCTGCTtttCATCATCAT |
| 5 | TGACGCAAACCTGCAGTTTCTCAGAAACTTTTCAGCATTTCA | TGACGCAAACCTGCAGTTTCTCAGAAACTTTTCAGCATTTCAtttCATCATCAT |
| 6 | TCGGTTCTCAAGTTCAGAGTTGCACTGCATGCTTACCATT | TCGGTTCTCAAGTTCAGAGTTGCACTGCATGCTTACCATTtttCATCATCAT |
| 7 | TAATCATTCAGTTGCCGTGCGAAGTATGGTCCATGTACTTCA | TAATCATTCAGTTGCCGTGCGAAGTATGGTCCATGTACTTCAtttCATCATCAT |
| 8 | GTCCGAGCAGGAGCTATAGTCGCTGGTTAGGCTCCCGTTATA | GTCCGAGCAGGAGCTATAGTCGCTGGTTAGGCTCCCGTTATAtttCATCATCAT |
| 9 | GCGTAGTAAAGTATGTCTGCGCCGGTAGATCCAGGATTTC | GCGTAGTAAAGTATGTCTGCGCCGGTAGATCCAGGATTTCtttCATCATCAT |
| 10 | CTGGAAATCAGCGGGCTTGCGATGGCATAGGATGGATACT | CTGGAAATCAGCGGGCTTGCGATGGCATAGGATGGATACTtttCATCATCAT |
| 11 | GTACTGCGGCGTCTGTAGCATAGACTGGGTTGTAGCGTAG | GTACTGCGGCGTCTGTAGCATAGACTGGGTTGTAGCGTAGtttCATCATCAT |
| 12 | AGATTCTCGTACTCTTGCGAATTCAGAATCTTCTGCGCCA | AGATTCTCGTACTCTTGCGAATTCAGAATCTTCTGCGCCAtttCATCATCAT |
| 13 | GTCAAACGTAATAAAGCCGTAACCTTTAGAAGCGCCTGTTCG | GTCAAACGTAATAAAGCCGTAACCTTTAGAAGCGCCTGTTCGtttCATCATCAT |
| 14 | AAAATTTCTAAATTCGTTTTCGTCGGCGTGGGGCGGTATT | AAAATTTCTAAATTCGTTTTCGTCGGCGTGGGCGGTATTtttCATCATCAT |
| 15 | CTCAGTCGCAGAGGATTTAGCCAGAAGACTCAGATCAGCA | CTCAGTCGCAGAGGATTTAGCCAGAAGACTCAGATCAGCAtttCATCATCAT |
| | ***Mlig455_028180 \| syt11 (Synaptotagmin) \| 2 \| p27 (tttCATCATCAT)*** | |
| 1 | ATTAGCGAATTATTTGTTGCTGTTTGTCCTCACGAGAGCA | ATTAGCGAATTATTTGTTGCTGTTTGTCCTCACGAGAGCAtttCATCATCAT |
| 2 | TTTTAGAAAACTCGCACGGGAACAGAAATTTCTTGCCA | TTTTAGAAAACTCGCACGGGAACAGAAATTTCTTGCCAtttCATCATCAT |
| 3 | GCTTTTAGTAATTCTCGCTTCAACGCTCGCGTCCTT | GCTTTTAGTAATTCTCGCTTCAACGCTCGCGTCCTTtttCATCATCAT |
| 4 | TTCCGCATCGGAGCGGCTATATCAAATAGGTCCATT | TTCCGCATCGGAGCGGCTATATCAAATAGGTCCATTtttCATCATCAT |
| 5 | GGGGCGGATCGACAGAAACACGAGAGATGATTCACA | GGGGCGGATCGACAGAAACACGAGAGATGATTCACAtttCATCATCAT |
| 6 | TCGACACAGACAAGGAGCGGCCACATCTAAGCCAAA | TCGACACAGACAAGGAGCGGCCACATCTAAGCCAAAtttCATCATCAT |
| 7 | AAGTTCAGTTATCACATTGACTGACGGATTCCGGGG | AAGTTCAGTTATCACATTGACTGACGGATTCCGGGGtttCATCATCAT |
| 8 | GGGCGAACGCCTCAAGTTCATGCGCTACACAACAAG | GGGCGAACGCCTCAAGTTCATGCGCTACACAACAAGtttCATCATCAT |
| 9 | AAAATAGACACGGCTACGGGAAATCTGGGAGAAGTT | AAAATAGACACGGCTACGGGAAATCTGGGAGAAGTTtttCATCATCAT |
| 10 | TTGCGCAAATAATTCTACATAAGCGCCTTTCCCGCG | TTGCGCAAATAATTCTACATAAGCGCCTTTCCCGCGtttCATCATCAT |
| 11 | TTTTAAGGACCCAAATACAGACATCGCAAGCGGATAGA | TTTTAAGGACCCAAATACAGACATCGCAAGCGGATAGAtttCATCATCAT |
| 12 | GAGGCTGCGGTTGTGCTAAGTGCGCGTTAACATAAA | GAGGCTGCGGTTGTGCTAAGTGCGCGTTAACATAAAtttCATCATCAT |
| 13 | AGTTCTTTAATTTCCAAGTTCCTTGATCTGACAAGCGGGG | AGTTCTTTAATTTCCAAGTTCCTTGATCTGACAAGCGGGGtttCATCATCAT |
| 14 | TTTTCAAACGGACGAAAATGACGGAACAGCGACGAA | TTTTCAAACGGACGAAAATGACGGAACAGCGACGAAtttCATCATCAT |
| 15 | CGCGAGCGACACGAATGACGAGGAGACAGAGTTGAG | CGCGAGCGACACGAATGACGAGGAGACAGAGTTGAGtttCATCATCAT |
| 16 | GAACGGAGAGGCCAGAATGAGGGCTGACAGAGACAG | GAACGGAGAGGCCAGAATGAGGGCTGACAGAGACAGtttCATCATCAT |
| 17 | GGGGCGTTAGAAATGAGAGCCAGATTCACTTGTCGG | GGGGCGTTAGAAATGAGAGCCAGATTCACTTGTCGGtttCATCATCAT |
| 18 | GATCACTTCGTTCTTGGTGACTCGGTCCCAGTCCAG | GATCACTTCGTTCTTGGTGACTCGGTCCCAGTCCAGtttCATCATCAT |

| | | |
|---|---|---|
| 19 | CGTTTGGCGGCACTTCAAACACGAACGATTCGTTGT | CGTTTGGCGGCACTTCAAACACGAACGATTCGTTGTtttCATCATCAT |
| 20 | GGATTCAGCGTGCGCTTCTTCACGTGGGTCTTCTTC | GGATTCAGCGTGCGCTTCTTCACGTGGGTCTTCTTCtttCATCATCAT |
| 21 | GATCCGCTGCCCGTTGTACTGCAGGTAGAGCTTCAC | GATCCGCTGCCCGTTGTACTGCAGGTAGAGCTTCACtttCATCATCAT |
| 22 | GGTTCTGGTTCTTCACCTTGAACTGCCTCGGCTGAA | GGTTCTGGTTCTTCACCTTGAACTGCCTCGGCTGAAtttCATCATCAT |
| 23 | CTGGTCACGCTCAGTAGGTCGTCCGTATCCGTCTCG | CTGGTCACGCTCAGTAGGTCGTCCGTATCCGTCTCGtttCATCATCAT |
| 24 | GATGATGTGGTCGCGCGAGAACCTGTCGAAGCTCAG | GATGATGTGGTCGCGCGAGAACCTGTCGAAGCTCAGtttCATCATCAT |
| 25 | GCATCCCGTAGAAGCTGAACGTCTCCTCGTACACAG | GCATCCCGTAGAAGCTGAACGTCTCCTCGTACACAGtttCATCATCAT |
| 26 | ACGATGGTCACCAGCAGAGTTTGCTTGCACTTGTCG | ACGATGGTCACCAGCAGAGTTTGCTTGCACTTGTCGtttCATCATCAT |
| 27 | CTGGTTGTGCAGCGAAAGGTTCCGGCTGATCTTCTG | CTGGTTGTGCAGCGAAAGGTTCCGGCTGATCTTCTGtttCATCATCAT |
| 28 | TGCCGCCCAGACTCACTGGTTGCATAGCAGAATCAG | TGCCGCCCAGACTCACTGGTTGCATAGCAGAATCAGtttCATCATCAT |
| 29 | CCGTGCATAGGCCGAATCACTGGCGCCATAATTTGT | CCGTGCATAGGCCGAATCACTGGCGCCATAATTTGTtttCATCATCAT |
| 30 | GGGCTCGCTCACTCACGGTGAGATCTCCAGAACATG | GGGCTCGCTCACTCACGGTGAGATCTCCAGAACATGtttCATCATCAT |
| 31 | AAAACTCGGCAACAAGATGTAAAAGATGTAAAAGATGCGGC | AAAACTCGGCAACAAGATGTAAAAGATGTAAAAGATGCGGCtttCATCATCAT |

***Mlig455_060907 | piwi | 8 | p28 (tttCAACTTAAC)***

| | | |
|---|---|---|
| 1 | AAAAGAACGCAGAGTATGAACAGCTTATTAAGACCGCACCGG | AAAAGAACGCAGAGTATGAACAGCTTATTAAGACCGCACCGGtttCAACTTAAC |
| 2 | AAAACCGGATGAAAACGAGAAAACGCTGGCACAACACAAC | AAAACCGGATGAAAACGAGAAAACGCTGGCACAACACAACtttCAACTTAAC |
| 3 | AAGAACCTTTCAAAGAACTCAAGCGGCGTACATCGGTAGT | AAGAACCTTTCAAAGAACTCAAGCGGCGTACATCGGTAGTtttCAACTTAAC |
| 4 | AATCAATCAACTCAAAGCACTCAACTGTCGCGATTGGCTC | AATCAATCAACTCAAAGCACTCAACTGTCGCGATTGGCTCtttCAACTTAAC |
| 5 | CAGTTAAAGTACAGGTGGGTCAGCTTGTACGTCAGCTGCT | CAGTTAAAGTACAGGTGGGTCAGCTTGTACGTCAGCTGCTtttCAACTTAAC |
| 6 | CGGCTTGATGTTGGCGTCATTCACGTCCTCAATCATGTTG | CGGCTTGATGTTGGCGTCATTCACGTCCTCAATCATGTTGtttCAACTTAAC |
| 7 | CGGAACATGCGAGAGCTGACCAGCTTCTTCACCACAATGG | CGGAACATGCGAGAGCTGACCAGCTTCTTCACCACAATGGtttCAACTTAAC |
| 8 | GTAGGCAGGTCGCCGTTCTTCTCTCTGAATTTCTGCAGGG | GTAGGCAGGTCGCCGTTCTTCTCTCTGAATTTCTGCAGGGtttCAACTTAAC |
| 9 | CTCCCTTTACGATGGGCGAGTACGAGTAGTACTGGGTGAA | CTCCCTTTACGATGGGCGAGTACGAGTAGTACTGGGTGAAtttCAACTTAAC |
| 10 | GTGATAAGTGTCCATCCCGACGATCATCGTGCGCTTCATC | GTGATAAGTGTCCATCCCGACGATCATCGTGCGCTTCATCtttCAACTTAAC |
| 11 | GGCTTGTTCACGTAGCACAGTCGCTTGATGGAGTCGTAGC | GGCTTGTTCACGTAGCACAGTCGCTTGATGGAGTCGTAGCtttCAACTTAAC |
| 12 | GAACACCAGGTCCAGGCCTTGGTCGATGTTTTCCTCAATC | GAACACCAGGTCCAGGCCTTGGTCGATGTTTTCCTCAATCtttCAACTTAAC |
| 13 | TCGTCTCGCTGGGAGTACACGAAGATCCAGTTCTTGCAGT | TCGTCTCGCTGGGAGTACACGAAGATCCAGTTCTTGCAGTtttCAACTTAAC |
| 14 | CATCTCATCAGTCAGACCGGTCATAAAGCAAACCTCTGGC | CATCTCATCAGTCAGACCGGTCATAAAGCAAACCTCTGGCtttCAACTTAAC |
| 15 | CAGAATCGGCTGCTCACTGTCCGACAAATTGACATTGTAC | CAGAATCGGCTGCTCACTGTCCGACAAATTGACATTGTACtttCAACTTAAC |
| 16 | AGCGAAATCTCCTCGCTCTGGCGAGTGCTCTTATTGTACC | AGCGAAATCTCCTCGCTCTGGCGAGTGCTCTTATTGTACCtttCAACTTAAC |
| 17 | TCGATTGCGTCAATACGATACGTCTTGTTGTTGTGCCGCG | TCGATTGCGTCAATACGATACGTCTTGTTGTTGTGCCGCGtttCAACTTAAC |
| 18 | CGGCTTGGAAATTGCCGGGATTGCGATTGTACATGTCGTA | CGGCTTGGAAATTGCCGGGATTGCGATTGTACATGTCGTAtttCAACTTAAC |
| 19 | AGAACCGTGCTCATGTGGATGATCTTGTGCGACACGTCAA | AGAACCGTGCTCATGTGGATGATCTTGTGCGACACGTCAAtttCAACTTAAC |
| 20 | CGATGCTGTGGAAGCTTCGTCGCTTGGTCGGGGTAATAAA | CGATGCTGTGGAAGCTTCGTCGCTTGGTCGGGGTAATAAAtttCAACTTAAC |
| 21 | ACCTGTCTAATTGCAACTTCGCCGCTCCACGACTTGATGC | ACCTGTCTAATTGCAACTTCGCCGCTCCACGACTTGATGCtttCAACTTAAC |
| 22 | CACCATTTGTCCTCGGTGTTGATCCGAACGAAATTCGTCA | CACCATTTGTCCTCGGTGTTGATCCGAACGAAATTCGTCAtttCAACTTAAC |
| 23 | CCGCGGCCTTTAGGCTGTTGAGACATGCTGAGTAACTTGG | CCGCGGCCTTTAGGCTGTTGAGACATGCTGAGTAACTTGGtttCAACTTAAC |

***Mlig455_034550 | tnnt2 (Troponin) | 2 | p28 (tttCAACTTAAC)***

| | | |
|---|---|---|
| 1 | TTTTCTCAGTCGGAGATTTGGGACAGTTGGCTAAGA | TTTTCTCAGTCGGAGATTTGGGACAGTTGGCTAAGAtttCAACTTAAC |
| 2 | TGCTTTTGGAAGCGCTAATGAGTTTCTGTCGTATTCCA | TGCTTTTGGAAGCGCTAATGAGTTTCTGTCGTATTCCAtttCAACTTAAC |
| 3 | TTTTGACTGACATTCTTCTGAGTGCATTTCCAGTGGTAGA | TTTTGACTGACATTCTTCTGAGTGCATTTCCAGTGGTAGAtttCAACTTAAC |
| 4 | AGAGAAAAGCGGGGAGGCACAAGCGGGAAATTTGTT | AGAGAAAAGCGGGGAGGCACAAGCGGGAAATTTGTTtttCAACTTAAC |
| 5 | CGATTTAAAAGTGAAGAGTTGGACATTGCTACTCGCAGACT | CGATTTAAAAGTGAAGAGTTGGACATTGCTACTCGCAGACTtttCAACTTAAC |
| 6 | GGAGAGTTGGCTGAATAATTTCAAAGATGCCCGGTT | GGAGAGTTGGCTGAATAATTTCAAAGATGCCCGGTTtttCAACTTAAC |
| 7 | GCGGAGGCCAATGAAGAAGAAGGACTGAACATTTAG | GCGGAGGCCAATGAAGAAGAAGGACTGAACATTTAGtttCAACTTAAC |
| 8 | AAATGAGCCTCCAGTCCGTTTAGAGGCGATTAAAGA | AAATGAGCCTCCAGTCCGTTTAGAGGCGATTAAAGAtttCAACTTAAC |
| 9 | AAAATTATTAAAGTCTTCCGTTGCTTTCAGCTTCACTCGGC | AAAATTATTAAAGTCTTCCGTTGCTTTCAGCTTCACTCGGCtttCAACTTAAC |
| 10 | TCTCAGTCAGGACGATCTTCGAGCTGGGCTTGATCC | TCTCAGTCAGGACGATCTTCGAGCTGGGCTTGATCCtttCAACTTAAC |
| 11 | GCGGTGGTCTTTGACTCGCTCGTACTGGCTGTACAT | GCGGTGGTCTTTGACTCGCTCGTACTGGCTGTACATtttCAACTTAAC |
| 12 | CAAGTTTCTCGGACAGCGGGTCAGGAGCAGTCTGAA | CAAGTTTCTCGGACAGCGGGTCAGGAGCAGTCTGAAtttCAACTTAAC |
| 13 | CGAGCGCGCTCTGTGAGCTCGACCATCTCCATATTC | CGAGCGCGCTCTGTGAGCTCGACCATCTCCATATTCtttCAACTTAAC |
| 14 | AGACTCGCTGAATCCGTCAATATTCAGCGGCTGAAC | AGACTCGCTGAATCCGTCAATATTCAGCGGCTGAACtttCAACTTAAC |
| 15 | CAGCCTCGCCGCCAGACTTCTTGCTGATTGTGAAAT | CAGCCTCGCCGCCAGACTTCTTGCTGATTGTGAAATtttCAACTTAAC |
| 16 | GCCTCGTTACGACCGCTTTCGTCTTCAGCAAAGCTG | GCCTCGTTACGACCGCTTTCGTCTTCAGCAAAGCTGtttCAACTTAAC |
| 17 | CTCCTCCTCGACTGTCGGCTCTTGGTTCTCTTCGTC | CTCCTCCTCGACTGTCGGCTCTTGGTTCTCTTCGTCtttCAACTTAAC |
| 18 | CGCTGTTGACAACTGGGCTCTGCCGACAAACAGAAA | CGCTGTTGACAACTGGGCTCTGCCGACAAACAGAAAtttCAACTTAAC |
| 19 | AGTTGATATTAAAGAAATGTTTTGCTCGGCTCGGCTGG | AGTTGATATTAAAGAAATGTTTTGCTCGGCTCGGCTGGtttCAACTTAAC |
| 20 | CGCCAGTAAGTGTCTGCGCATTTATTCCCATTCCTG | CGCCAGTAAGTGTCTGCGCATTTATTCCCATTCCTGtttCAACTTAAC |

***Mlig455_022240 | apob (Lipoprotein amino terminal region) | 2 | p30 (tttAATACTCTC)***

| | | |
|---|---|---|
| 1 | TCAGAAGAAAGAGATCGCCCTAGGCTCAGAGTCGATCGAG | TCAGAAGAAAGAGATCGCCCTAGGCTCAGAGTCGATCGAGtttAATACTCTC |
| 2 | GATGATGATGCGCCTGTCTGAGGTGATCTTCACGGTAACA | GATGATGATGCGCCTGTCTGAGGTGATCTTCACGGTAACAtttAATACTCTC |
| 3 | CTGATCTCGAACTTCACTAGGTGCTCCTTGGCGTCAATGT | CTGATCTCGAACTTCACTAGGTGCTCCTTGGCGTCAATGTtttAATACTCTC |
| 4 | AAACCTTGTTGTGGTAGAACGGAGAGACGCGGATCTCAAC | AAACCTTGTTGTGGTAGAACGGAGAGACGCGGATCTCAACtttAATACTCTC |
| 5 | GAACGCCAGATGTACTTCCTGTTTGGACCGGTAACGTTCA | GAACGCCAGATGTACTTCCTGTTTGGACCGGTAACGTTCAtttAATACTCTC |
| 6 | CAGACAGTTCGGAGTCAACCAGAAGCTTCTTCAGCTCAGA | CAGACAGTTCGGAGTCAACCAGAAGCTTCTTCAGCTCAGAtttAATACTCTC |
| 7 | GATCTCGATGGTAGCGATTTCCCTCTCGTCATCGTTGATG | GATCTCGATGGTAGCGATTTCCCTCTCGTCATCGTTGATGtttAATACTCTC |
| 8 | ATGTGATCCTCTCAGAGCTTAGGTCCTTCGGGGTAAAAGC | ATGTGATCCTCTCAGAGCTTAGGTCCTTCGGGGTAAAAGCtttAATACTCTC |
| 9 | CTCCAGCTTGATCTGGGGCACCTTCTTAGAATCGTCTTCC | CTCCAGCTTGATCTGGGGCACCTTCTTAGAATCGTCTTCCtttAATACTCTC |
| 10 | CAGCGACTCCGAGATGGTAGAGAAAGCGTCGTTCTCAAAC | CAGCGACTCCGAGATGGTAGAGAAAGCGTCGTTCTCAAACtttAATACTCTC |
| 11 | GAGGACATCCTTGTTGACCTTCAAAGAGGTTAGCCACTGG | GAGGACATCCTTGTTGACCTTCAAAGAGGTTAGCCACTGGtttAATACTCTC |
| 12 | CGCGAGTGTTCTTCATGTAGACGATGTCGTAGCCCATCAG | CGCGAGTGTTCTTCATGTAGACGATGTCGTAGCCCATCAGtttAATACTCTC |
| 13 | GGAATCCTTGACGACTTCGAGCATCGTCTGGAGCATCTTC | GGAATCCTTGACGACTTCGAGCATCGTCTGGAGCATCTTCtttAATACTCTC |
| 14 | GTGGTAGACGTTGAGATTGGCACGAGTGACAAAGTCCCTC | GTGGTAGACGTTGAGATTGGCACGAGTGACAAAGTCCCTCtttAATACTCTC |
| 15 | GCTCCTCAACGATTTCGTTCAGATCCTGCTCCTTGTTCAG | GCTCCTCAACGATTTCGTTCAGATCCTGCTCCTTGTTCAGtttAATACTCTC |
| 16 | CCTCATTAGAGTCGATCTTGAGCAGCTGGTCGAACAGCTG | CCTCATTAGAGTCGATCTTGAGCAGCTGGTCGAACAGCTGtttAATACTCTC |

***Mlig455_025291 | pcna (Proliferating cell nuclear antigen) | 2 | p30 (tttAATACTCTC)***

| | | |
|---|---|---|
| 1 | ACAAAATGCGCCATAAGCTGGACAAATTAACTACTTGATCAGC | ACAAAATGCGCCATAAGCTGGACAAATTAACTACTTGATCAGCtttAATACTCTC |
| 2 | AAAATAGACAGCTCGTCATTACAGTCAGTGCATATGGTGCA | AAAATAGACAGCTCGTCATTACAGTCAGTGCATATGGTGCAtttAATACTCTC |
| 3 | TATGGTGCAGATGCACTGGATGAAGCGTGCCCTAGCA | TATGGTGCAGATGCACTGGATGAAGCGTGCCCTAGCAtttAATACTCTC |
| 4 | CGCTGTACCGTAGAGATTCGTCATCAACAATAATTATCGTGCA | CGCTGTACCGTAGAGATTCGTCATCAACAATAATTATCGTGCAtttAATACTCTC |
| 5 | AAGTAAAACTACTTGTCAGCTTCGTCCTCCATCTTGGGAG | AAGTAAAACTACTTGTCAGCTTCGTCCTCCATCTTGGGAGtttAATACTCTC |
| 6 | CGGATATAGCCCAGATCGGCAATATTGTACTCAATCAGAGC | CGGATATAGCCCAGATCGGCAATATTGTACTCAATCAGAGCtttAATACTCTC |
| 7 | TGCCGCCTTGCTGAAGATGTTGAAGTAGCGCAGCGAGTA | TGCCGCCTTGCTGAAGATGTTGAAGTAGCGCAGCGAGTAtttAATACTCTC |
| 8 | CAGCCGAAGGCATCCTGATCACAGCCTTGTAGTCGGTTT | CAGCCGAAGGCATCCTGATCACAGCCTTGTAGTCGGTTTtttAATACTCTC |
| 9 | ATGCCCAGATGCTCAGCATCCAAGTCCATCAGCTTCA | ATGCCCAGATGCTCAGCATCCAAGTCCATCAGCTTCAtttAATACTCTC |
| 10 | TACTCAGACACTTTCTCTTGGTTCGGCATCGACTCGA | TACTCAGACACTTTCTCTTGGTTCGGCATCGACTCGAtttAATACTCTC |
| 11 | GTGATGGCATCCGAGTTGCTGGCGCACTTCAGAATCT | GTGATGGCATCCGAGTTGCTGGCGCACTTCAGAATCTtttAATACTCTC |
| 12 | CAGCTCTGCACGTTCATGCCTAGCGACACGTTTCTGT | CAGCTCTGCACGTTCATGCCTAGCGACACGTTTCTGTtttAATACTCTC |
| 13 | CTGTAAGTCTCAAATCCCTCAGACTTGAGCAGCATTGAGA | CTGTAAGTCTCAAATCCCTCAGACTTGAGCAGCATTGAGAtttAATACTCTC |
| 14 | CTAACATGACTGCTGTCCATCGCCTGCAGACTGATGC | CTAACATGACTGCTGTCCATCGCCTGCAGACTGATGCtttAATACTCTC |
| 15 | CAGGTGGCTTCGTTAACCAGCTCGCGTAGAGCCTCAATCA | CAGGTGGCTTCGTTAACCAGCTCGCGTAGAGCCTCAATCAtttAATACTCTC |
| 16 | TTCAGGTACTCGCCCTGGACAAGCTTGGCCTCAAACA | TTCAGGTACTCGCCCTGGACAAGCTTGGCCTCAAACAtttAATACTCTC |

***Mlig455_017161 | sperm1 (M. lignano testes-specific gene) | 2 | p30 (tttAATACTCTC)***

| | | |
|---|---|---|
| 1 | ACGCCATGGCAAGAATATTCATAGTTTCGATTCGAGCTGGT | ACGCCATGGCAAGAATATTCATAGTTTCGATTCGAGCTGGTtttAATACTCTC |
| 2 | AGAGAGCCGCAATTCAGTAGTCAACTTTCAACGCACTTTT | AGAGAGCCGCAATTCAGTAGTCAACTTTCAACGCACTTTTtttAATACTCTC |
| 3 | AAAATGCTTTGAAAACAAAATTCGACAGCCAGTTTGGGGAGG | AAAATGCTTTGAAAACAAAATTCGACAGCCAGTTTGGGGAGGtttAATACTCTC |
| 4 | AAAAGCGGCAGACTTCCCCTGAAAGCAGATAGAAATACCA | AAAAGCGGCAGACTTCCCCTGAAAGCAGATAGAAATACCAtttAATACTCTC |
| 5 | TTAAATTAATGACGAGTAGAGTCTGTGGCCTGGACCTCGC | TTAAATTAATGACGAGTAGAGTCTGTGGCCTGGACCTCGCtttAATACTCTC |
| 6 | CGAAGAGCGGCTCGCTTTCCACCTTCCTTGACTTGAGCAT | CGAAGAGCGGCTCGCTTTCCACCTTCCTTGACTTGAGCATtttAATACTCTC |
| 7 | GGGTACACGTGAGACATCACCGTCTCCATGCGGTCATTCT | GGGTACACGTGAGACATCACCGTCTCCATGCGGTCATTCTtttAATACTCTC |
| 8 | TGTCCACGATGAGATACTTTAGCCAGCGGTAGCGGCCCAT | TGTCCACGATGAGATACTTTAGCCAGCGGTAGCGGCCCATtttAATACTCTC |
| 9 | AGTCACCTCGATTTCGCACTTGGTCCGGTTTGCAACGGAC | AGTCACCTCGATTTCGCACTTGGTCCGGTTTGCAACGGACtttAATACTCTC |
| 10 | CGGGATCGACTAAGAATGCCTGAATCATCTCGTCCGACAG | CGGGATCGACTAAGAATGCCTGAATCATCTCGTCCGACAGtttAATACTCTC |
| 11 | TAGCTTTCTACGCGTCTCCCTGGCTTCGTCATCTTTTCCT | TAGCTTTCTACGCGTCTCCCTGGCTTCGTCATCTTTTCCTtttAATACTCTC |
| 12 | GCCGATGTCTTCAGAGGCGCCGCTCTGATCTAGAGACTCATA | GCCGATGTCTTCAGAGGCGCCGCTCTGATCTAGAGACTCATAtttAATACTCTC |
| 13 | CAGTGCGTTAGACAAGTTGGAGTCGCTGTCCATGATAATGT | CAGTGCGTTAGACAAGTTGGAGTCGCTGTCCATGATAATGTtttAATACTCTC |
| 14 | GTTGCTGTTGTTTCCCGCTCTGATCACCCGCAGCATTTGC | GTTGCTGTTGTTTCCCGCTCTGATCACCCGCAGCATTTGCtttAATACTCTC |
| 15 | TTGCTGCTGCTTCTCTGGCTGTCTTTCGATTGCATGTTTT | TTGCTGCTGCTTCTCTGGCTGTCTTTCGATTGCATGTTTTtttAATACTCTC |
| 16 | CAGTTGTTGGACGTCTGTCGGAACTGGCGCTTGCAGTGTA | CAGTTGTTGGACGTCTGTCGGAACTGGCGCTTGCAGTGTAtttAATACTCTC |
| 17 | TGACGAGCTCAATCTGTCTGAGTGGATTCTGGCTTTCAGC | TGACGAGCTCAATCTGTCTGAGTGGATTCTGGCTTTCAGCtttAATACTCTC |

**Table S3.** Primer Exchange Reaction (PER) results on the ordered probe oligo pools and probes' pool performance in AP SABER

| Target gene | Approximate probe pool length after probe PER concatemerization (bp) | Concentration after the probe pool after column purification (ng/ul) | Expression pattern in AP SABER | Signal development time [first stopped, last stopped] (minutes) |
|---|---|---|---|---|
| *boll* | 280 | 148 | Testes, ovaries - meiotic cells | [10, 40] |
| *syt11* | 300 | 178 | Neural system: brain and peripheral | [33, 75] |
| *piwi* | 300 | 109 | Stem cells and gonads | [38, 60] |
| *tnnt2* | 300 | 90 | Muscles | [42, 46] |
| *apob* | 220 | 54 | Gut and phagocytes | [14, 21] |
| *pcna* | 200 | 58 | Stem cells and gonads | [115, 146] |
| *sperm1* | 170 | 50 | Testes | [11, 56] |

**Table S4.** Costs of the key reagents for the signal development reactions

| Development Protocol (conditions) | Entry costs (Euro)* | Number of reactions* | Limiting factors | Costs/reaction (Euro) |
|---|---|---|---|---|
| direct SABER FISH (3 targets) | 720 | 540 | labeled oligonucleotides | 1,33 |
| AP SABER (1 target, FAM, NBT/BCIP) | 610 | 3540 | DIG-labeled oligo | 0,17 |
| *SABER TSA (2 targets, DIG+FAM, 2 tyramides)* | 860 | 610 | Fluorescent tyramides | 1,41 |
| pSABER TSA (2 targets, 2 tyramides) | 4200 | 610 | Fluorescent tyramides | 6,89 |
| SABER HCR (3 targets, all adapters) | 1071 | 600** | Labeled hairpin amplifiers | 1.79** |

** see Table S5 for more details and costs' justification*

*** with 3x re-use of the amplification mix*

**Table S5.** Detailed costs of the development reagents

| Application | Reagent | Producer | (nmol) Purchased | Stock Concentration | (µl) stock solution | (µl) stock/100 µl devel. solution | (Euro) total stock entry price | Euro/µl | Number of 1x reactions |
|---|---|---|---|---|---|---|---|---|---|
| direct SABER FISH | p27-SABER-ATTO633 | IDT | 5,4 | 10 µM | 540 | 1 | 240 | 0,44 | 540 |
| | p28-SABER-ATTO550 | IDT | 5,4 | 10 µM | 540 | 1 | 240 | 0,44 | 540 |
| | p30-SABER-ATTO488 | IDT | 5,4 | 10 µM | 540 | 1 | 240 | 0,44 | 540 |
| SABER HCR | p27-SABER_to_B1-HCR | IDT | 20 | 10 µM | 2000 | 1 | 19 | 0,01 | 2000 |
| | p28-SABER_to_B1-HCR | IDT | 20 | 10 µM | 2000 | 1 | 19 | 0,01 | 2000 |
| | p30-SABER_to_B1-HCR | IDT | 20 | 10 µM | 2000 | 1 | 19 | 0,01 | 2000 |
| | p27-SABER_to_B2-HCR | IDT | 20 | 10 µM | 2000 | 1 | 19 | 0,01 | 2000 |
| | p28-SABER_to_B2-HCR | IDT | 20 | 10 µM | 2000 | 1 | 19 | 0,01 | 2000 |
| | p30-SABER_to_B2-HCR | IDT | 20 | 10 µM | 2000 | 1 | 19 | 0,01 | 2000 |
| | p27-SABER_to_B3-HCR | IDT | 20 | 10 µM | 2000 | 1 | 19 | 0,01 | 2000 |
| | p28-SABER_to_B3-HCR | IDT | 20 | 10 µM | 2000 | 1 | 19 | 0,01 | 2000 |
| | p30-SABER_to_B3-HCR | IDT | 20 | 10 µM | 2000 | 1 | 19 | 0,01 | 2000 |
| | Pair of HCR hairpins B1* | Molecular Instruments | 0,6 | 3 µM | 200 | 1 | 300 | 1,50 | 200 |
| | Pair of HCR hairpins B2* | Molecular Instruments | 0,6 | 3 µM | 200 | 1 | 300 | 1,50 | 200 |
| | Pair of HCR hairpins B3* | Molecular Instruments | 0,6 | 3 µM | 200 | 1 | 300 | 1,50 | 200 |
| AP SABER/SABER TSA | p27-SABER-DIG | IDT | 35,4 | 10 µM | 3540 | 1 | 164 | 0,05 | 3540 |
| | p28-SABER-6FAM | IDT | 220 | 10 µM | 22000 | 1 | 77 | 0,00 | 22000 |
| | p30-SABER-6FAM | IDT | 220 | 10 µM | 22000 | 1 | 77 | 0,00 | 22000 |
| pSABER TSA | p27-SABER-5'HRP | Eurogentec | 60 | 10 µM | 6000 | 1 | 1800 | 0,30 | 6000 |
| | p28-SABER-5'HRP | Eurogentec | 60 | 10 µM | 6000 | 1 | 1800 | 0,30 | 6000 |
| SABER TSA | Anti-DIG-POD/HRP | Roche | - | 2000x | 200 | 0,05 | 317 | 1,59 | 4000 |
| | Anti-Fluorescein-POD/HRP | Roche | - | 2000x | 200 | 0,05 | 302 | 1,51 | 4000 |
| (p)SABER TSA | FITC Tyramide** | Biotium | 610 | 5 mM (500x) | 122 | 0,2 | 300 | 2,46 | 610 |
| | CF568** | Biotium | 610 | 5 mM (500x) | 122 | 0,2 | 300 | 2,46 | 610 |
| | CF647 Tyramide** | Biotium | 610 | 5 mM (500x) | 122 | 0,2 | 300 | 2,46 | 610 |
| AP SABER | Anti-DIG-AP | IDT | - | 2000x | 200 | 0,05 | 307 | 1,54 | 4000 |
| | Anti-Fluorescein-AP | IDT | - | 2000x | 200 | 0,05 | 362 | 1,81 | 4000 |
| | Vector Blue | Vector Labs | - | 110x | 3000 | 0,9 | 185 | 0,06 | 3333 |
| | NBT/BCIP | Roche | - | 50x | 8000 | 2 | 171 | 0,02 | 4000 |

**Amplification mix of HCR hairpins can be reused at least 3 times*

***Fluorescent tyramides can be synthesized in bulk for lesser costs* (Lauter et al., 2011)

**Table S6.** Strengths, weaknesses, and suggested best application for different OneSABER signal development protocols

**SABER protocol | development time | multiplexing capacity | 1x reaction costs (Euros) | signal**

| Advantages | Disadvantages | Suggested application |
|---|---|---|
| ***Direct SABER FISH \| 1 day for 3+ targets \| Excellent \| 1.33 \| fluorescent*** | | |
| • The fastest possible protocol.<br>• No antibodies, or intermediate amplification steps required.<br>• High permeability of secondary probes. Non-diffused, high-resolution signal. | • Low signal strength without branching or ordering more primary probes.<br>• High entry costs for purified fluorophore-labeled oligonucleotides compared to AP SABER and SABER TSA. | Multiplex FISH for strong and verified targets |
| ***AP SABER \| 1 day for 1 target \| Poor \| 0.17 \| colorimetric & fluorescent*** | | |
| • Signal development can be monitored in real time.<br>• The highest possible sensitivity due to long-lasting activity of the AP enzyme.<br>• High contrast signal due to absence of autofluorescence issue.<br>• AP fluorescent substrates do not bleach-out and can be used for high intensity confocal scans.<br>• Indefinite preservation of the signal and storage of the samples after mounting at room temperature.<br>• Can be easily combined with SABER TSA for dual multiplexing without significant changes in workflow. | • Majorly limited to single gene colorimetric *in situs*.<br>• Requires use of antibodies.<br>• For fluorescent substrates, color precipitates are relatively diffused, generate shadows and can mask other fluorophores if overdeveloped (the issue is not that prominent if using high magnification and high aperture objectives with confocal microscopy). | • Single gene *in situ*.<br>• Establishment of the ground-truth expression pattern and estimation of the gene expression strength (can help to select best FISH SABER approach).<br>• Optimization of the hybridization conditions prior to moving to other SABER protocols.<br>• Detection of weak and extremely weakly expressed genes. |
| ***SABER TSA \| 1 day per target, up to 3 targets \| Good \| 1.41 \| fluorescent*** | | |
| • Possibly the strongest fluorescent signal amplification technique due to combination of antibody and TSA enhancement.<br>• Can be easily combined with AP SABER for dual multiplexing without significant changes in workflow. | • Compared to pSABER and SABER HCR, different hapten-labeled (DIG/FAM/DNP) imager probes as well as corresponding anti-DIG/FAM/DNP HRP-conjugated antibodies are required.<br>• Multiplexing is time consuming with 1-2 extra days compared to pSABER TSA and SABER HCR.<br>• Compared to SABER HCR, the fluorescent signal is more diffused within a cell. | • Single gene or multiplex FISH when signal sensitivity is the priority.<br>• The most sensitive FISH protocol suitable for detection of weakly expressed genes. |
| ***pSABER TSA \| 1-2 days for up to 3 targets \| Good \| 6.89 \| fluorescent*** | | |
| • No antibodies and hapten-labeled oligonucleotides required, hence, faster implementation than with antibody-mediated SABER TSA.<br>• Otherwise, shares similar advantages to SABER TSA | • HRP-conjugated oligonucleotides are very costly as a first-time investment.<br>• Multiplexing is sequential and still more hands on time is required compared to SABER HCR.<br>• Compared to SABER HCR, the fluorescent signal is more diffused within a cell. | The same as SABER TSA, but when time is a higher priority. |
| ***SABER HCR \| 1 day for 3+ targets \| Excellent \| 1.79 \| fluorescent*** | | |
| • Fast SABER protocol, only 1 day slower than direct SABER FISH for all targets.<br>• One-step no-protein multiplexing and signal amplification.<br>• High permeability and low background of the HCR hairpins compared to HRP-labeled oligos and anti-hapten antibodies.<br>• Non-diffused, high-resolution signal.<br>• HCR amplification mix can be re-used at least 3 times. | • The cost of fluorophore-labeled HCR hairpins is high.<br>• Official HCR hairpins are distributed by only one company.<br>• Slightly more complicated experiment planning when it comes to combining HCR hairpins to different adapter oligonucleotides and starting sequences of pre-amplifying concatemers of the SABER probes. | Multiplex FISH with 2-4 moderately or strongly expressed genes when signal specificity and working time are the priorities. |

** see Tables S5 and S6 for justification and more details*

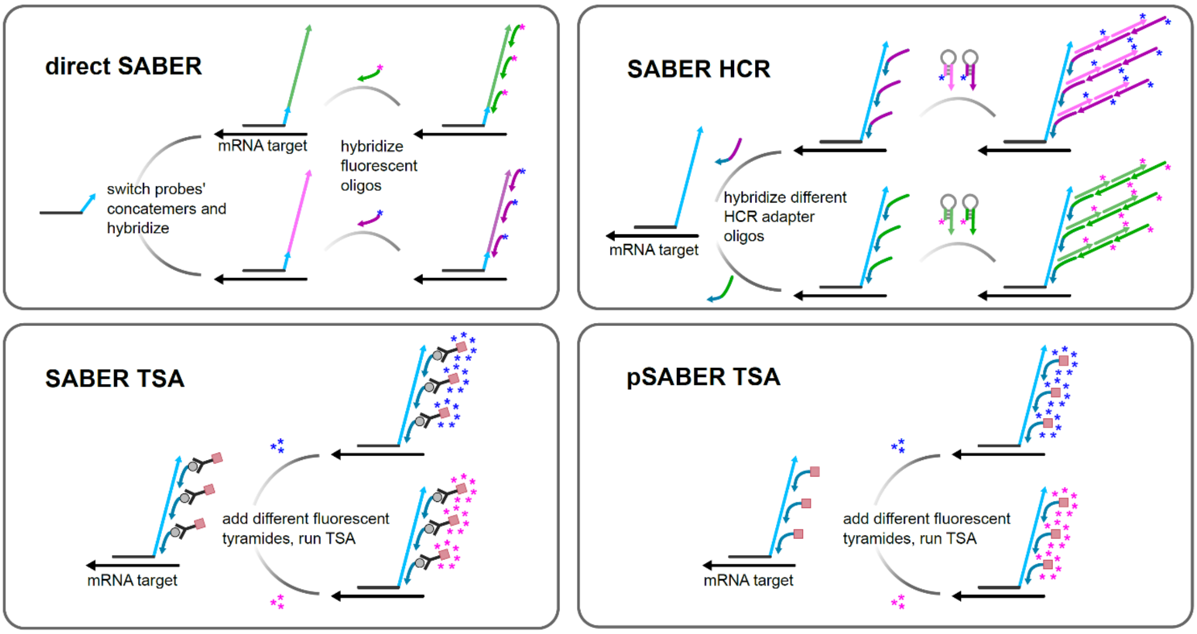


**Fig. S1. Schematics of fluorophore switching for different OneSABER FISH signal development methods.**

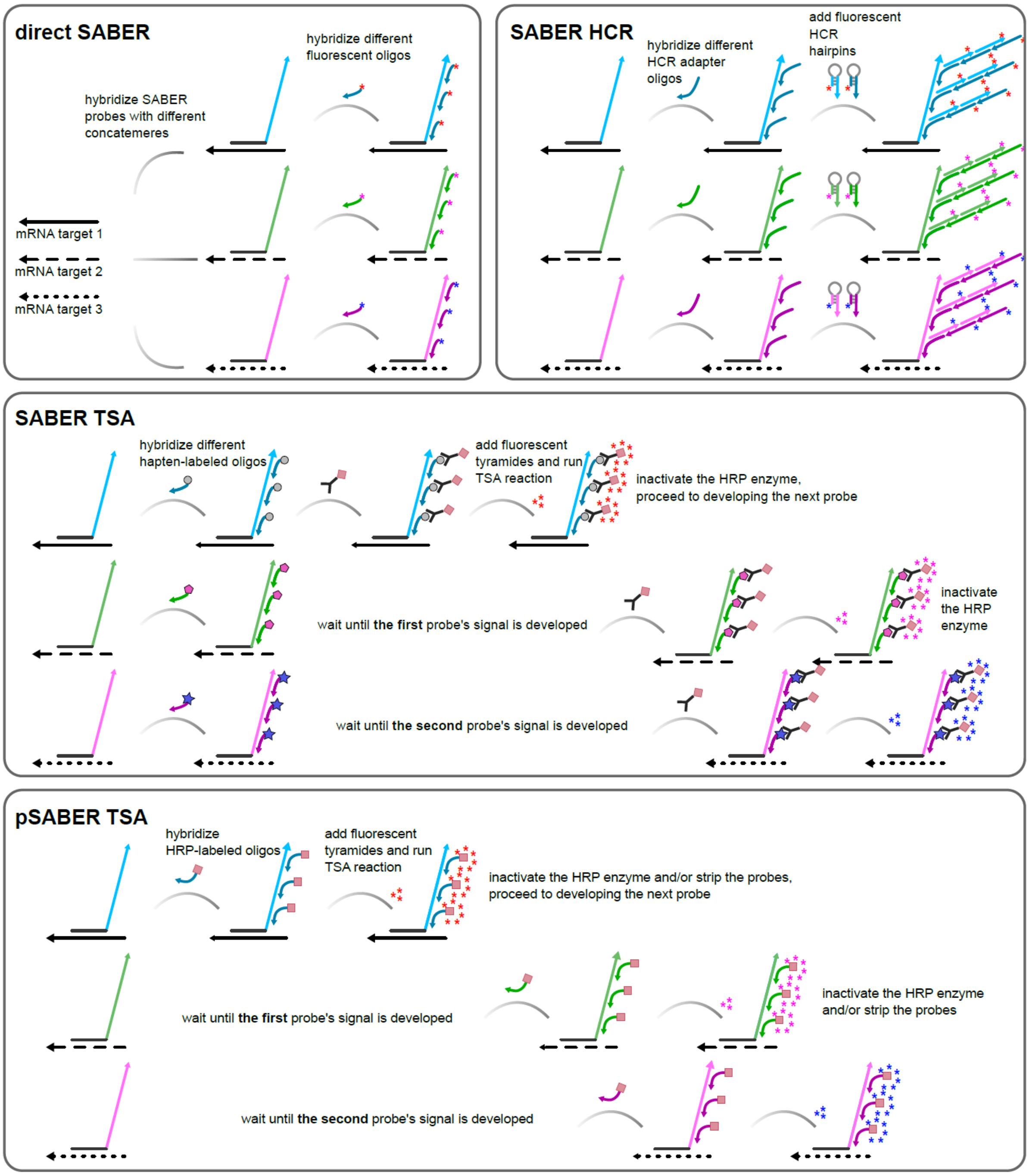


**Fig. S2. Schematics of multiplexing steps for different OneSABER FISH signal development methods.**

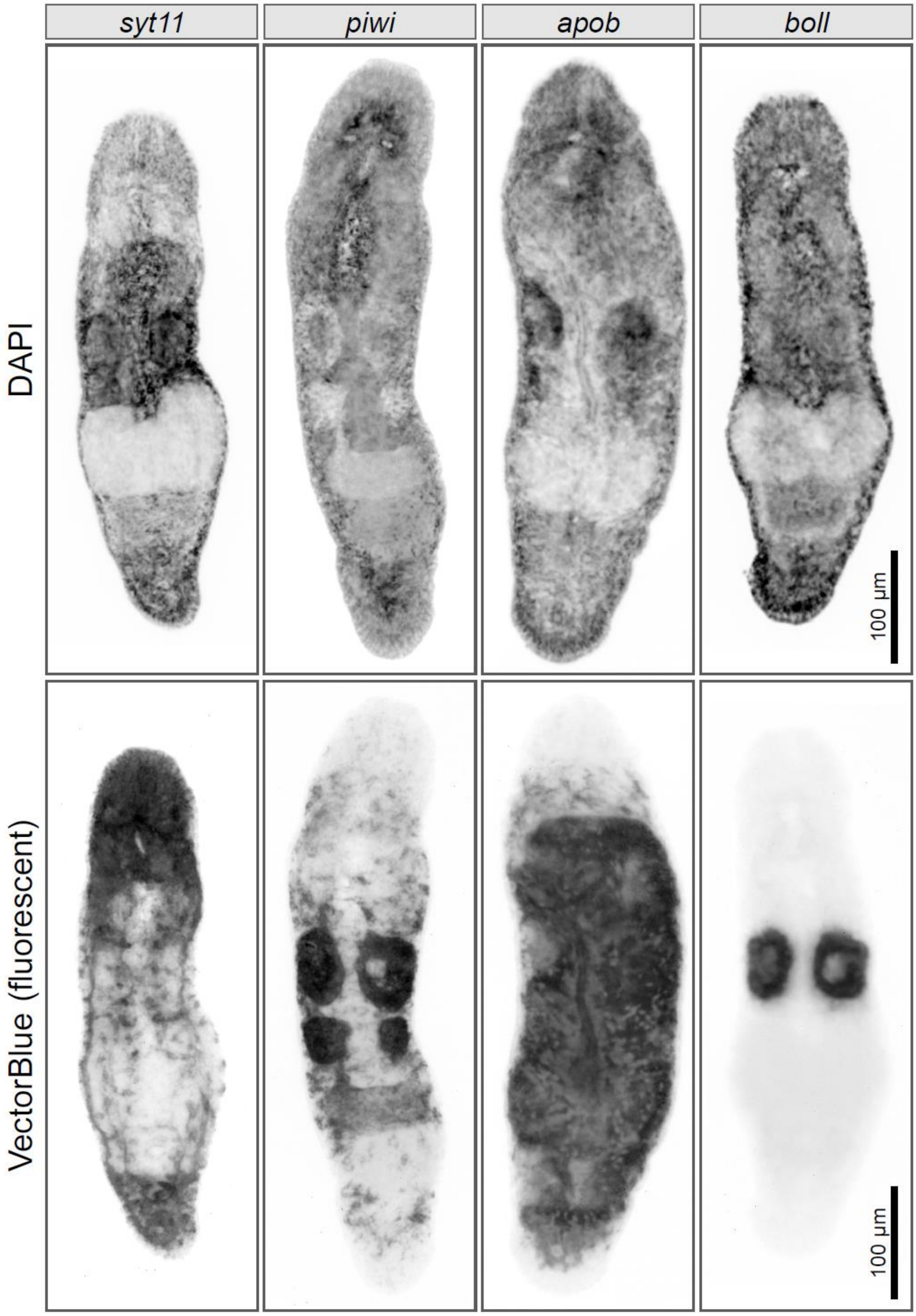


**Fig. S3. Vector Blue can be visualized fluorescently in near far-red spectrum.** Inverted images of widefield grayscale fluorescent photos of homeostatic *M. lignano* worms are shown. Note how the colorimetric signal masks the DAPI fluorescence.

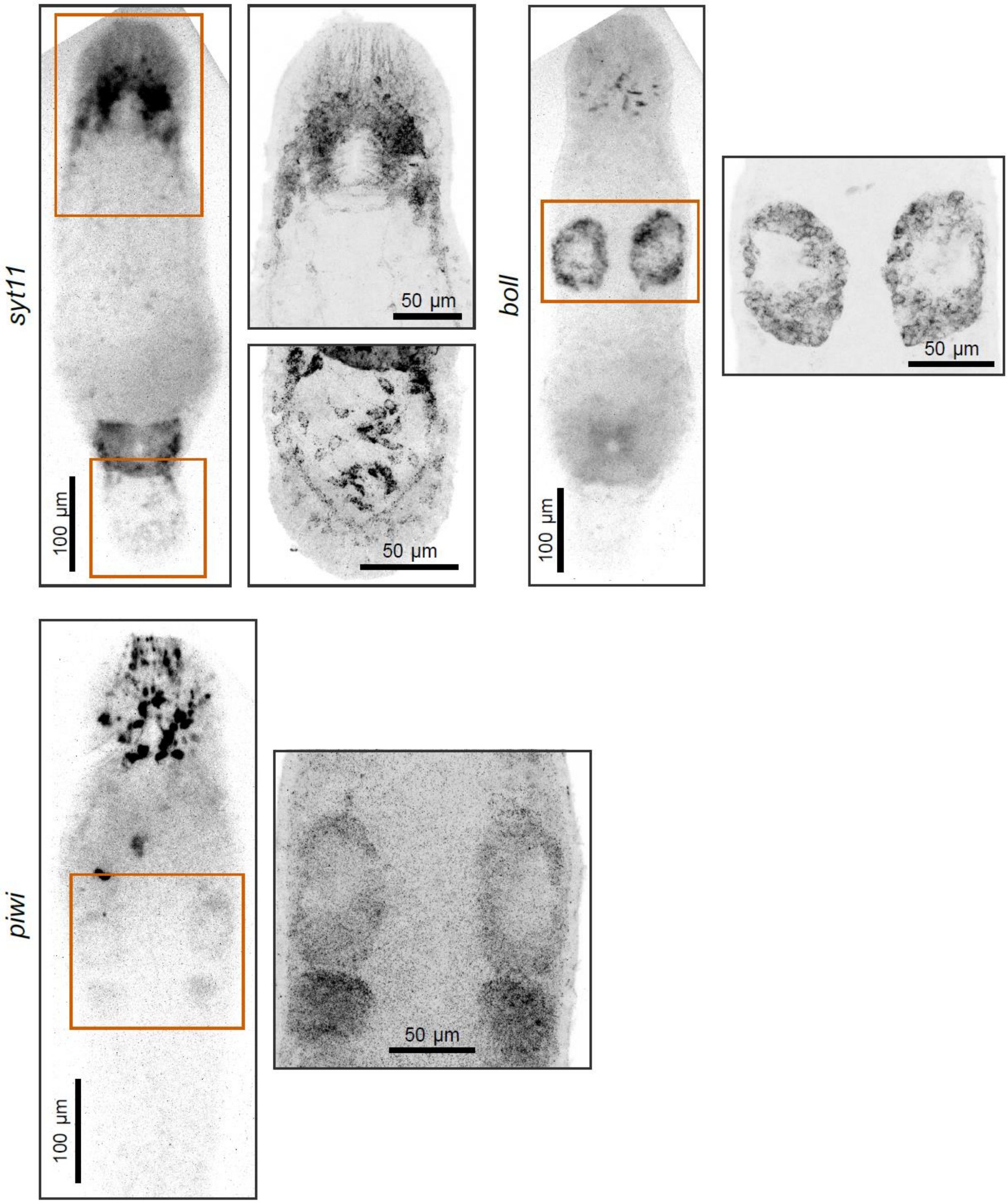


**Fig. S4. Direct SABER FISH with ATTO633 labeled secondary probes.** Inverted images of widefield grayscale fluorescent photos of homeostatic *M. lignano* worms are shown. Exposure time for the widefield photos for the same channel was ~17 times longer than on Figs 3, 4. Zoom-in regions are maximum intensity projections of several confocal microscopy slices.

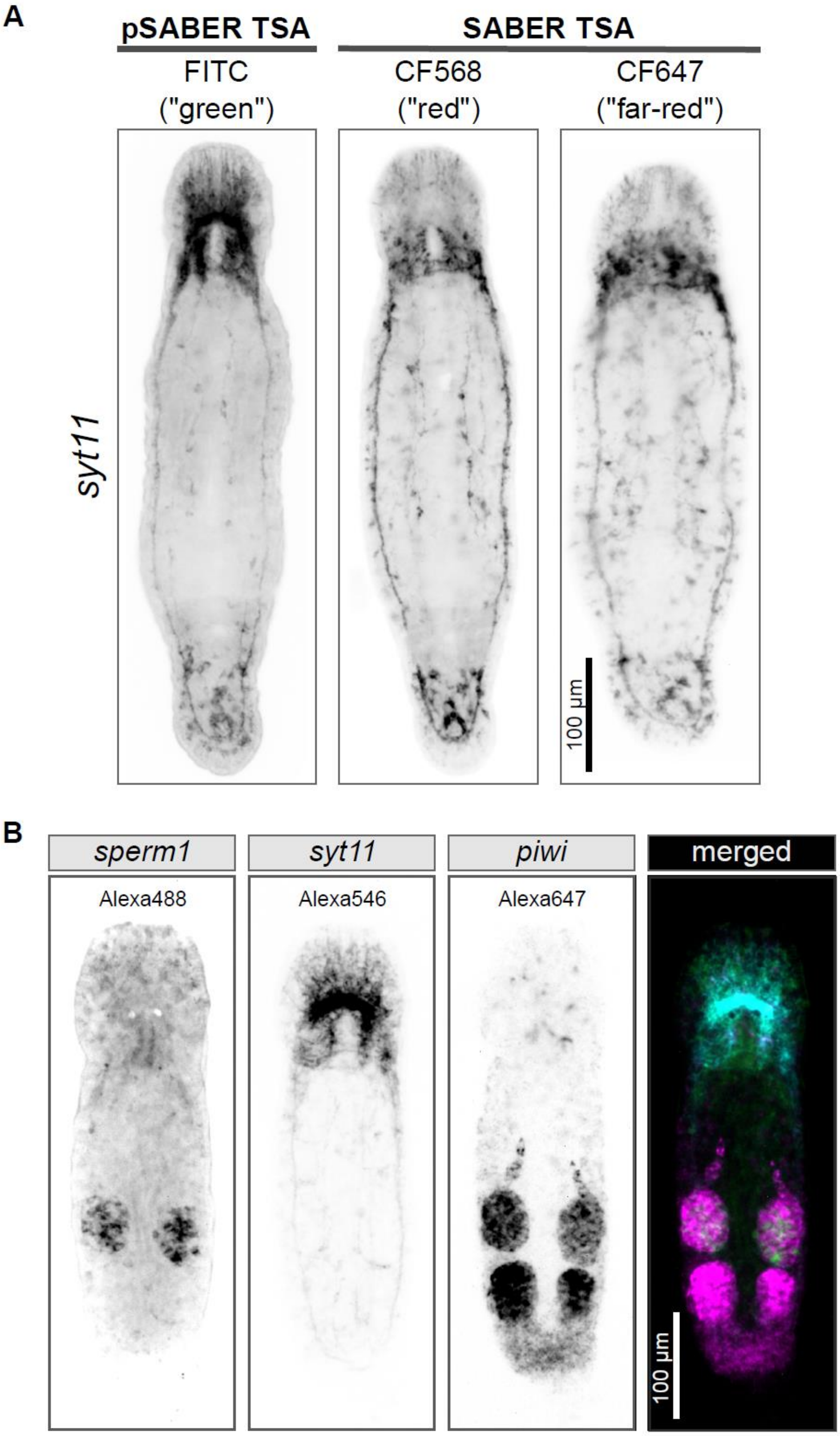


**Fig. S5. OneSABER TSA and HCR results after fluorophores' switching.** (A) Comparison between pSABER TSA and SABER TSA with different fluorescent tyramides. (B) SABER HCR after using different SABER-to-HCR adapters than on Fig. 4.

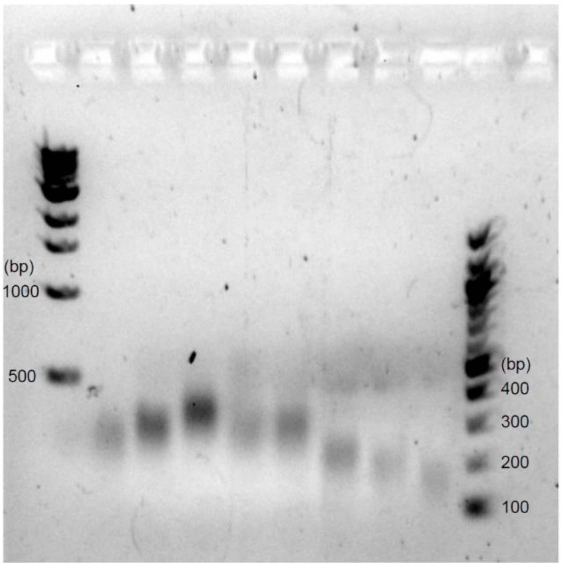


**Fig. S6. An example of a successful primer exchange reaction for concatemerization of different primary SABER probe pools.** The gel was prepared as in the Supplementary information Section S1.

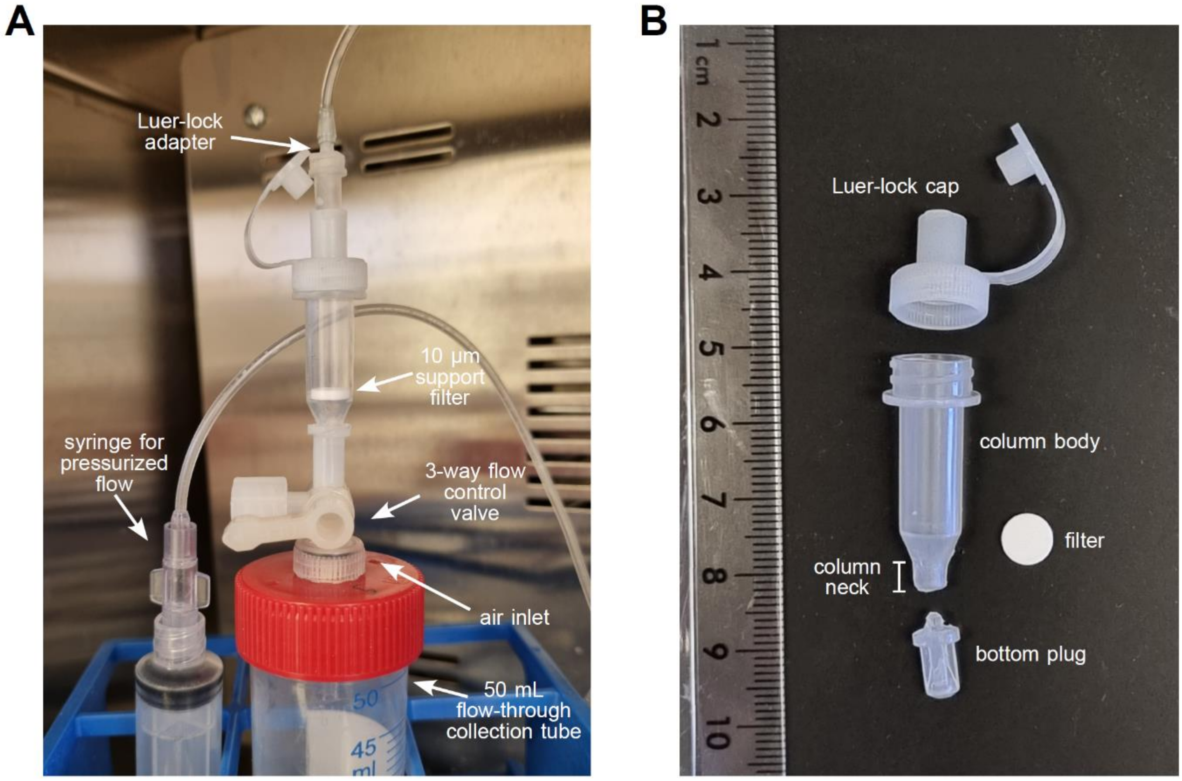


**Fig. S7. Illustration of the liquid exchange column assembly.** See Supplementary information Section S2 for more details.

### Section S1. Design, preparation, and validation of SABER probes

For more general and detailed information on the preparation of SABER probes, please refer to the original manuscript and its supplementary protocol (Kishi et al., 2019).

Here, we provide an adapted and verified version of the probe design, validation, and preparation for *Macrostomum lignano*. All steps were performed using Linux-based operating systems and tested on Ubuntu Desktop 21.04, but it should be easily adaptable to MacOS or Windows systems.

#### *Mining of SABER oligonucleotide probes*

Before you start, be sure to have Python v. 2.7 installed. It will be necessary to run the OligoMiner oligonucleotide search software. Download OligoMiner scripts (https://github.com/beliveau-lab/OligoMiner).

1) Obtain the spliced mRNA/cDNA sequence of your gene of interest (GOI). Make a reverse complement of it, using e.g. https://www.bioinformatics.org/sms/rev_comp.html or your favorite standalone sequence editing and annotation software (SnapGene, Benchling, UGENE, CLC workbench, etc.)). Save the reverse complement sequence in a FASTA-formatted text file.

2) Create an executable text file `OligoMinerScript.sh` with the following code (for Windows, an executable `*.bat` file should be created instead):

```
python [YOUR-PATH-TO-OLIGOMINER]/OligoMiner/blockParse.py\
                                            --minLength 40\
                                            --maxLength 43\
                                            --min_GC 40\
                                            --max_GC 60\
                                            --min_Tm 43\
                                            --max_Tm 48\
                                            --salt 390\
                                            --formamide 50\
                                            --Spacing 15\
                                            -b\
                                            --file $1

python ./bedToFasta.py
```

3) In the folder of the `OligoMinerScript.sh` Create a text file: `bedToFasta.py` with the following code:

```
import sys
args = sys.argv
if len(args) < 2:
print("Specify input BED file as the argument\n")
sys.exit(1)
outf = open(args[1]+'.fasta', 'w')
with open(args[1], 'r') as in_file:
```

```
for l in in_file:
l = l.split()
probe_id = "_".join(l[0:3])
probe_seq = l[3]
outf.write(">%s\n%s\n" % (probe_id, probe_seq))
outf.close()
```

4) Substitute `[YOUR-PATH-TO-OLIGOMINER]` to a relevant absolute path where you saved the OligoMiner scripts. Alternatively, add the OligoMiner folder to your system `$PATH` variable and directly call `blockParse.py`. We recommend saving and individually modifying a copy of the `OligoMinerScript.sh` along with `bedToFasta.py` for each GOI target in a separate folder for more efficient data management.

5) Parameters that should remain constant are indicated in bold, and they were optimized to the hybridization conditions used in our protocol (Supplementary Information, Section S4). The rest should be optimized and manually modified based on the desired number of probes if the program returns too little, or too many identified targets. Follow these recommendations:

a) `--maxLength` parameter `<= 48` and `--minLength >= 35`. If you get too many probes, narrow the min/max length gap, or broaden it if you get too little.
b) `--min_GC >= 30` and `--max_GC <= 65`. Narrow the GC gap for too many returned probes and broaden it otherwise.
c) `--min_Tm >= 42` and `--max_Tm <= 50`. Narrow the Tm gap for too many returned probes and broaden it otherwise.
d) `--Spacing >= 5`. Increase spacing between probes when get too many probes.

6) Open a command line/Terminal app. Navigate to the folder with the GOI reverse complement FASTA file (`GOI_rev_comp_seq.fasta`) and a copy of `OligoMinerScript.sh`.

7) From the terminal, execute the following line of code:

```
sh ./OligoMinerScript.sh GOI_rev_comp_seq.fasta
```

8) In the terminal, the script should return an identified number of probes that fit the specified stringency criteria. In the working folder two following files should be created:

```
GOI_rev_comp_seq.bed
GOI_rev_comp_seq.bed.fasta
```

9) The `*.bed` file is easy to copy into a table/spreadsheet software like Excel or LibreOffice Calc, while the `*.fasta` file can be readily used as an input into BLAT/BLAST software to verify and QC the matches.

10) If you get way too many predicted probe binding sites, modify `OligoMinerScript.sh` to match more stringent criteria as recommended above.

11) Once you are close to the desired number of probes, proceed to their QC.

12) For *M. lignano*, we use BLAT software of the genome browser ([https://gb.macgenome.org/cgi-bin/hgBlat?hgsid=29881_ae6Xo7vwVUbknBvVRAFgCkZ2pbGg&command=start](https://gb.macgenome.org/cgi-bin/hgBlat?hgsid=29881_ae6Xo7vwVUbknBvVRAFgCkZ2pbGg&command=start)) to check for the on- and off-targets of the predicted probe set on the transcriptome and the genome of the worm by copying and pasting the predicted probe sequences from the generated `*.bed.fasta` file.

13) A good probe should:

a) Perfectly match (100% identity, 100% sequence span) only found once specifically to the "negative" strand of the GOI or a known number of times if various low-divergent paralogs or pseudogenes are present, with the number of matches equal to the number of paralogs/pseudogenes.
b) Do not significantly align to a repetitive part of your GOI and any other mRNA targets.

14) Remove the probes that do not pass the criteria from the `*.bed` file, then, to update the FASTA-file in the terminal execute separately:

```
python ./bedToFasta.py
```

If you got too little probes left, relax the selection parameters and re-execute the script:

```
sh ./OligoMinerScript.sh GOI_rev_comp_seq.fasta
```

15) Once you get the desired number of verified probes, add the same 12 base pair Primer Exchange Reaction (PER) 3' initiator sequence to the 3' ends of each probe. Use:

a) "`tttCATCATCAT`" for the p27 primer;
b) "`tttCAACTTAAC`" for the p28 primer;
c) "`tttAATACTCTC`" for the p30 primer.

More diverse PER primers, although untested in this article, can be found in the original SABER study Supplementary materials (Kishi et al., 2019).

16) Proceed to ordering the designed sequences as desalted dry oligonucleotides in a 96-well plate format to save the costs. All the oligonucleotides used in this study were ordered from Integrated DNA Technologies (IDT, Table S2).

***How many probes is enough?***

While hybridization specificity is determined by the probes' design. The sensitivity of the *in situ* hybridization is directly proportional to the number of probes. However, a good number of probes should provide a balance between sensitivity and costs. We suggest dividing the genes by their transcriptional expression into 3 categories: strongly, medium, and weakly expressed. The expression levels can be known a priori from previous studies or known RNA-seq profiles of tissues, organs, and/or whole organisms. As a rule of thumb, we suggest, if it is allowed by the length of the target, using ~10-15 probes for strongly expressed genes, 15-20 probes for medium expressed genes and 20+ probes for weakly expressed genes. Since later more probes can always be ordered, and sensitivity of the assay can be adjusted by the probe concatemerization length, we suggest testing the protocol starting from 15 probes per gene.

***Dilution and preparation of stock SABER oligonucleotide mixes***

Upon arrival, dissolve each probe oligonucleotide to 100 µM with $Ca^{2+}$/$Mg^{2+}$-free 1xPBS or Tris-water (pH ~7-8). This will be your long-term storage stocks. For each GOI probe set, combine equal volumes (e.g. 10 µL) of the dissolved probes in one tube making a GOI 100 µM stock probe pool. Make a 10 µM dilution of the pool. This is your ready-to-use oligo pool for *in vitro* concatemerization by PER reaction.

***Setting up concatemerization of SABER oligonucleotides by PER reaction in vitro***

Assembly of a PER reaction is according to the original SABER FISH protocol (Kishi et al., 2019). Mix all the following components on ice in a 200 µL PCR tube/strip, excluding the probe oligo pool, and adding the Bst polymerase last.

| **Component** | **Volume (µL, total volume = 100 µL)** |
|---|---|
| $Ca^{2+}$/$Mg^{2+}$-free 10xPBS | 10 |
| 100 mM MgSO4* (NEB) | 10 |
| dNTP mix* (Only A, C, and T, 6mM each, NEB) | 5 |
| Clean.G oligonucleotide (1 µM) * | 10 |
| Bst DNA Polymerase, Large Fragment (8 U/µL, NEB) | 2.5 |
| PER Hairpin (5 µM) *, ** | 10 |
| Mili-Q $H_2O$ | 42.5 |
| Probe oligo pool (10 µM) | 10 |

** Oligonucleotide sequences and their function are described in Table S1*

*** the PER hairpin must match the 3' initiator sequence of the probes in oligo pool*

Pre-incubate the mix in a thermocycler for 15 min at 37°C, then add the probe oligo pool, mix well, and return to the thermocycler for a desired extension time at 37°C. The extension time will determine the length of the concatemers and will vary for different PER hairpins. An optimal extension time should be determined empirically and will vary from application to application. Although long probes can provide higher sensitivity, they can also give stronger background and cause non-specific binding due to lower penetration in whole-mount specimens. We found that probes of 300-500 bp in length provide a reasonable balance between sensitivity and signal-to-noise ratio. We suggest starting with a 1 h extension and then adjusting the length of the reaction by 15 min.

Stop the extension by heat inactivation of the enzyme at 80°C for 20 min., cool the reaction down to 4°C.

***Assessing the quality and length of PER probe concatemerization and SABER probes purification***

Here, we propose an alternative to the original protocol and a much more sensitive method to estimate the length of the single-stranded DNA concatemers by gel electrophoreses. Prepare a 1% agarose gel in TAE/TBE buffer with added Ethidium Bromide (the cheapest option) or other DNA staining dye. Take 1 μL of the heat inactivated PER concatemerization reaction and mix it with 1 μL of 10 μM matching unlabeled universal "dummy" antisense oligonucleotide secondary probe (p27 to p27, p28 to p28, p30 to p30, see Table S1) and 8 μL of 1xPBS in a PCR tube. Anneal the secondary probe to the concatemer by heating up to 95°C for 1 min, cooling down to 37°C, incubating for 1 min, and cooling down to 4°C. Mix with your preferred gel loading dye, load on the gel, load 100 bp dsDNA ladder marker and run the gel for 30-40 min at 80-90V or until desired separation is achieved. Analyze the gel. A good direct PER reaction should result in a seemingly single-size distinct band on the gel with, potentially, some smearing (Fig. S6).

In this method, we avoid the problem of very low sensitivity of DNA binding dye to ssDNA by converting its major part to dsDNA by annealing cheap unlabeled oligonucleotides. Additionally, only 1 μL of the PER reaction is used for the analysis instead of 10 μL recommended by the original SABER protocol (Kishi et al., 2019).

If the reaction was successful, proceed to the direct column purification for the remaining 99 μL of the mix. We use Qiagen PB binding buffer at 7:1 buffer-to-sample ratio to efficiently bind short (200-500 bp) ssDNA. The mixture is then loaded into gel/PCR purification silica columns (Favorgen, Germany) according to the manufacturer's protocol. The elution is done in 40-60 μL of 1xPBS. The final ssDNA concentration and yield was estimated by NanoDrop 2000. We find that a yield of 40-120 ng/μL is acceptable and 1-2 μL of the probes can be used for 1 round of *in situ* hybridization.

***Setting up switching PER concatemerization reaction***

Let's consider an example of switching from p27 to p28 concatemer sequence.

| **Component** | **Volume (μL, total volume = 100 μL)** |
|---|---|
| $Ca^{2+}/Mg^{2+}$-free 10xPBS | 10 |
| 100 mM MgSO4* (NEB) | 10 |
| dNTP mix* (Only A, C, and T, 6mM each, NEB) | 5 |
| Clean.G oligonucleotide (1 μM)* | 10 |

| | |
|---|---|
| Bst DNA Polymerase, Large Fragment (8 U/µL, NEB) | 2.5 |
| p28 PER Hairpin (5 µM) * | 10 |
| Switch p27 to p28 PER Hairpin (5 µM)* | 1 |
| Mili-Q $H_2O$ | 38.5 |
| Probe oligo pool (10 µM) | 10 |

** Oligonucleotide sequences and their function are described in Table S1*

Follow the same procedure as with the direct PER extension. Extension time and the resulting length of concatemers will change and may require additional optimization. The bands after switch reaction may appear less sharp on the gel, although without noticeable effect on *in situ* hybridization.

### Section S2. Assembly and operation of hybridization columns

The idea of using *in situ* mini-columns to facilitate fast liquid exchange, increase reproducibility, and minimize sample loss was borrowed from the article of S.Q. Irvine (Irvine, 2007). Column assembly and operation is largely the same as described in the paper. However, we introduced the following modifications (Fig. S7):

1) changed 15 mL flow-through collection tubes to 50 mL collection tubes;

2) removed small narrow point 10 μm filters to allow unrestricted gravity flow;

3) introduced flow control valves for easier and unpressurized liquid exchange;

A column in our design consists of a column body, a main support 10 μm filter, and a Luer-lock adapter screw cap. Columns are attached through a flow control 2/3-way valve to a 50 mL flow-through collection tube (Fig. S7A). Liquid exchange is performed either passively, by just opening the cap and the valve, or actively, by applying additional pressure through a syringe.

To ensure lick-proof sealing, the column neck (Fig. S7B) is gently scraped by sandpaper or scalpel, and then lightly lubricated with Vaseline petrol jelly. For storage and/or handling of columns with liquid and samples outside the assembly, a removable bottom plug can be used (Fig. S7B)

Each column can be reused more than 20 times without noticeable loss of performance. Between each *in situ* experiments columns are disassembled and their parts thoroughly washed in mili-Q water and soaked overnight. To remove any residual probes, DNA, RNA, and protein contaminants, the parts are soaked in 1M NaOH solution for 10 min and then thoroughly washed in mili-Q. We did not notice any cross-contamination or substantial loss of sensitivity/specificity of ISH between the experiments. After washing columns are left to air-dry on clean paper tissues and then pre-assembled and stored in a closed but air-ventilated dust-free container at room temperature until necessary.

### Section S3. Recipes for the *in situ* hybridization protocol in *M. lignano*

**Stock solutions** better prepared in advance. Store all stock solutions at room temperature, if not specified otherwise.

7.14% $MgCl_2$*6H2O (100 mL):

- Dissolve 7.14 g of MgCl2*6$H_2O$ in Mili-Q water (adjust the volume to 100 mL). Pass through 0.22-0.45 µm filter;

1xPBSTw (50 mL):

- 5 mL of 10xPBS (calcium and magnesium-free);
- 500 µL of 10% Tween 20 (0.1%);
- Mili-Q water up to 50 mL;

HCR amplification buffer (5xSSCTw, 50 mL):

- 12.5 mL of 20xSSC;
- 500 µL of 10% Tween 20 (0.1%);
- Mili-Q water up to 50 mL;

2xSSCTw (50 mL):

- 5 mL of 20xSSC;
- 500 µL of 10% Tween 20 (0.1%);
- Mili-Q water up to 50 mL;

WashHyb (500 mL, store in 15 mL tubes at -20°C, consumption 3 mL per column):

- 50 mL of 20xSSC (2x);
- 200 mL of 100% deionized formamide (40%);
- 25 mL of 10% Tween 20 (0.5%);
- 225 mL of Mili-Q water;

PreHyb (150 mL, store in 2 mL tubes at -20°C, consumption 250 µL per column):

- 15 mL of 20xSSC (2x);
- 60 mL of 100% deionized formamide (40%);
- 3 mL of 50xDenhard's solution (1x);
- 3 mL of 10 mg/mL salmon sperm DNA (200 µg/mL) (1x);
- 1.5 mL of 10 mg/mL Heparin (100 µg/mL);
- 15 mL of 10% Tween 20 (1%);
- 52.5 mL of Mili-Q water;

Hyb (100 mL, store in 2 mL tubes at -20°C, consumption 170 µL per column):

- 10 mL of 20xSSC (2x);
- 40 mL of 100% deionized formamide (40%);
- 2 mL of 50xDenhard's solution (1x);
- 2 mL of 10 mg/mL salmon sperm DNA (200 µg/mL) (1x);
- 1 mL of 10 mg/mL Heparin (100 µg/mL);
- 10 mL of 10% Tween 20 (1%);
- 20 mL of 50% (w/v) Dextran sulphate in Mili-Q water (10%);
- 20 mL of Mili-Q water;

Vector Blue buffer (10 mL):

- 9.9 mL of 0.1 M Tris-HCl (pH 8.5);
- 100 μL of 10% Tween 20 (0.1%);

NBT/BCIP buffer, aka NTMT (10 mL):

- 5 mL of 0.2M Tris-HCl pH 9.5 (0.12M);
- 1 mL of 1 M NaCl (0.1 M);
- 0.5 mL of 1 M $MgCl_2$ (50 mM);
- 0.5 mL 10% Tween 20 (0.5%);
- 3 mL of Mili-Q water;

After mixing, let the buffer sit for 5-10 min at room temperature to allow potential salt crystals to form, and then pass the solution through a 0.22-0.45 μm syringe filter.

Glycerol mounting solution (50 mL):

- Weigh 31.7 g of 100% glycerol (80%);
- 50 μL of 10% Tween 20 (0.01%);
- Mili-Q water to 50 mL;

Sodium Azide stop solution (15 mL, store ar 4°C):

- 98 mg of Sodium Azide;
- PBSTw up to 15 mL;

**Solution to always prepare fresh right before use**

*General solutions*

4% formaldehyde fixation solution (2 mL):

- 267 μL of 30% formaldehyde (MeOH-free);
- 1733 μL of 1xPBS (no Tween!!!);

4% formaldehyde re-fixing solution (1 mL):

- 134 μL of 30% formaldehyde (MeOH-free);
- 866 μL of 1xPBSTw;

ProteinaseK permeabilization solution (2 mL):

- 2 μL of 20 mg/mL proteinaseK stock solution (1/1000 dilution, stored at –20°C);
- 2 mL of PBSTw;

$H_2O_2$/Formamide Bleaching and permeabilization solution (2 mL):

- 1750 μL of Mili-Q water;
- 100 μL of 100% Deionized Formamide (5%, stored at –20°C);
- 50 μL of 20xSSC (0.5x);
- 80 μL of 30% $H_2O_2$ (1.2%);
- 20 μL of 10% Tween 20 (0.1%);

*Alkaline phosphatase colorimetric ISH development substrate solutions*

Vector Blue development solution (2.5 mL):

- 2.5 mL of Vector Blue buffer (prepared in advance);
- 1 drop (~40 μL) of Vector Blue Reagent 1;
- 1 drop (~40 μL) of Vector Blue Reagent 2;
- 1 drop (~22.5 μL) of Reagent 3;

Add Reagents 1-3 sequentially, mixing well in-between.

NBT/BCIP development solution (2 mL):

- 1960 μL of filtered NBT/BCIP buffer (NTMT, prepared in advance);
- 40 μL of NBT/BCIP stock solution (Roche, stored at 4°C);

*Tyramide Signal Amplification (TSA) FISH buffers*

TSA antibody blocking buffer (2 mL):

- 100 μL of Horse Serum (5%);
- 100 μL of Roche Western Blocking Reagent (RWBR, 5%);
- 1800 μL 1xPBSTw;

TSA amplification buffer (2 mL):

- 1 μL of 200 mg/ml 4-iodophenol (in 100% EtOH, 1/2000 dilution, stored at 4°C);
- 20 μL of 0.3% $H_2O_2$ (prepare fresh from 30% $H_2O_2$ by 1/100 dilution in 1xPBS);
- 1979 μL 1xPBS;

TSA development buffer (500 μL):

- 0.5-2 μL of 5 mM fluorescent tyramide DMSO stock (stored at –20°C);
- 499 μL of freshly made TSA FISH amplification buffer;

### Section S4. *In situ* hybridization protocol in *M. lignano*

#### Before you start, mind that if not specified otherwise…

1) All transfers of worms are done using a P200 pipette with low-retention pipette tips.
2) Up to 400 worms can be fixed simultaneously using the protocol.
3) All steps are done at ambient room temperature.
4) Wash volume for columns is ***400 µL***, and 500 µL for 2 mL tubes.
5) All the specified washing durations as well as the number of washing steps are the minimum recommended and surely can be increased, being only beneficial for the signal/noise ratio.

### S4.1 Day (0-1). Worms' preparation and fixation

***Starvation***

0) Transfer worms for at least ***48h*** in a Petri dish with ***20 mL*** of ASW without diatoms. After the first ***24h***, transfer the worms again to a new Petri dish with fresh ASW.

***Note:*** *accommodate starvation time for your regeneration experiment. For example, if you perform a* ***24h*** *regeneration experiment, worms can be cut* ***24h*** *after starvation, and continue fasting after amputation for an additional* ***24h***.

***Relaxation and immobilization***

1) Transfer the starved worms in a drop of ASW on the lid of the starvation Petri-dish. Using light, allow them to move and concentrate in a corner of the drop.

2) Using as little pipetting steps as possible, transfer the worms to the bottom corner of a small Petri dish (keep the lid separate for step (4)). Remove as much excessive ASW as possible taking care of not drying out any worm.

3) Add ***3 mL*** of ***7.14% $MgCl_2$*6H2O*** to the worms. Immediately start swirling the dish to allow even mixing. Be sure that all the worms are detached from the bottom and from each other. Once all the worms are immobile, proceed to the next step.

- ***Notes:*** *if worm tails attach to the bottom, use gentle pipetting near the worms without sucking them up in the tip until all animals are detached.*
- *Successful relaxation will result in slightly bent "banana-shape" worms, passively hovering due to cilia beating or completely immobile.*
- *Different worm ages, size, and conditions require different relaxation time which can vary from* ***3 – 8 min***. *Do not leave them longer than* ***10 mins*** *in* ***$MgCl_2$ solution***.

***Fixation and 1st permeabilization***

4) Collect the worms in the center of the dish by swirling using gentle circular motion. Using as little as possible pipetting steps, transfer the immobilized worms to the center of the saved small Petri dish from step (2). While gently vibrating and swirling the lid, aspirate as much as possible of the ***$MgCl_2$ solution***.

5) Using a P1000 pipette, sequentially add ***2 mL*** of ***4% formaldehyde*** fixative solution (1 mL + 1 mL) while vibrating/shaking the lid to allow even mixing. Start your timer set to ***30 min***.

6) Approximately ***15 sec*** later, while shaking, add drop by drop ***200 µL*** of ***100% glacial acetic acid***.

- **Note:** *addition of acetic acid may cause some worms to stick to the bottom of the lid. Use gentle pipetting near the worms (avoid sucking them in!) to detach them.*

7) After ***25 min*** of fixation, add ***2 µL*** of ***10% Tween*** and mix it well by pipetting. Start transferring the worms from the lid into a 2 mL microcentrifuge tube.

8) After ***30 min*** of fixation, remove the fixative and perform ***4x 5 min*** washes with ***500 µL*** of ***PBSTw***.

- **Note:** *in between washing steps, allow worms to sink down. Take care not to aspirate worms and always leave some liquid behind to avoid drying of the worms. Use a stereomicroscope to assist in the process.*
- **Critical note:** *for still unknown reasons (potentially addition of acetic acid), after 1-2 washes in* ***PBSTw*** *worms form clumps and clusters and stick to each other. Most of the clusters can be separated by passing them through a P200 tip (small clusters) or P1000 tip (big clusters). You may still attempt to separate most of the worms in* ***50% MeOH/PBSTw*** *and* ***100% MeOH*** *during steps (9) and (10). Once separated, worms do not stick to each other again.*

***Dehydration and storage***

9) Perform one buffer exchange and one 5 min wash with 50% MeOH/PBSTw, and then repeat the same with ***100% MeOH***.

10) Keep the fixed worms in ***100% MeOH*** for at least ***30 min*** if you want to proceed with hybridization on the same day. Alternatively, store them at **–20°C** for several months until needed.

### S4.2 Day 1. 2nd Permeabilization and hybridization of primary SABER probes

#### *Rehydration*

1) Transfer fixed worms stored in ***100% MeOH*** from **–20°C** to room temperature ***100% MeOH***.

- **Note:** *if some worms remain stuck to each other and pipetting does not help, you must separate them mechanically using, e.g. a soft synthetic brush bristle glued to the tip of a Pasteur glass pipette.*

2) Perform one buffer exchange and one ***5 min*** wash with ***50% MeOH/PBSTw***. Wash ***3x 5 min*** with ***PBSTw***.

#### 2nd permeabilization

3) pre-heat freshly made ***ProteinaseK solution*** to **37°C**. Remove PBSTw from the worms and add ***500 µL*** of the pre-heated ***ProteinaseK solution***. Depending on age of worms, their size, and state (regenerating/homeostatic) use the following treatment times at **37°C** as a starting guideline:

| ***Worms' age/stage*** | ***ProteinaseK treatment time*** |
|---|---|
| Hatchlings/amputated heads | Omit, proceed directly to step (6) |
| Juveniles/Tail-regenerating adults/1-2-week-old adults | 10 min |
| 3–4-week-old adults | 14 min |

4) One minute before the end of the incubation time, move the tube to room temperature, remove ***ProteinaseK*** solution and briefly, but delicately wash once with ***1 mL*** of ***PBSTw***.

5) Remove PBSTw and add ***500 µL*** of freshly made ***4% formaldehyde re-fixing solution***. Incubate for ***15 min*** at room temperature. Remove the fixative and wash ***3x 5 min*** with ***2xSSCTw***.

6) Prepare fresh ***2 mL of $H_2O_2$/formamide bleaching solution***. Remove ***2xSSCTw*** and add the bleaching solution to the worms. Incubate for at least ***45 min*** to ***1 h*** under direct bright white LED light illumination.

At the same time, assemble hybridization column(s) and take out ***preHyb*** (***250 µL***/column), ***Hyb*** (***170 µL***/column), and ***WashHyb*** (***3 mL***/column) buffer aliquots from **-20°C**.

7) Wash worms ***3x 5 min*** in ***2xSSCTw***.

***Pre-hybridization and hybridization***

8) Move the worms from the 2 mL tube into a small Petri dish filled with ***~3 mL*** of ***2xSSCTw***.

9) Move the desired number of worms (up to 30 adults) into hybridization column(s) loaded with ***400 µL 2xSSCTw***. Perform one buffer exchange and ***2x*** consecutive ***5 min*** washes with ***WashHyb***. Move the columns, ***preHyb***, ***Hyb***, and the remaining ***WashHyb*** buffer to a preheated to **55ºC** hybridization oven.

- **Note:** *from this point all the steps onward are performed with the samples/columns inside the oven. The remaining **WashHyb** will be used on the next day for post-hybridization washes.*

10) Exchange ***WashHyb*** to ***250 µL***/column of preHyb and incubate for at least ***30 min***. Prepare SABER probe hybridization solution by adding ***1-1.5 µL*** of dedicated SABER probe(s) per ***170 µL***/column of ***Hyb*** buffer and mix well. Keep the solution at **55ºC** until the end of pre-hybridization.

11) Exchange ***preHyb*** to ***Hyb***. Decrease the oven temperature to **48ºC**. Leave the sample hybridizing ***overnight*** or for ***16+ hours***.

- **Note:** ***48ºC*** *is a good starting hybridization temperature for most of the probes. However, some probes may require **2ºC** higher temperatures if signal is strong but non-specific background is also noticeable, or lower if no signal is detected. In the same manner, lowering the probes concentration may be beneficial for better signal/noise ratio.*

### S4.3 Day 2. Post-hybridization washes, hybridization of secondary probes, and signal development

#### *Post-hybridization washes*

1) Add ***250 µL*** of warm ***WashHyb*** buffer on top of the ***Hyb*** solution and then slowly flush the column.

2) Do ***1x*** brief ***WashHyb*** buffer exchange, and then ***2x 30 min*** washes with ***400 µL WashHyb***. Put ***2xSSCTw*** to warm up to **48°C** in the oven.

3) Do ***1x*** brief buffer exchange to ***2xSSCTw*** and then ***2x 5 min*** washes with ***2xSSCTw***.

- **Note:** *pause point. If necessary, samples can be stored for **1-2 days** at **4°C**.*

4) Cool the oven to **42°C** and pre-warm ***PBSTw***. Keep ***PBSTw*** at **42°C** until the antibody blocking steps.

5) Do ***1x*** buffer exchange and then ***2x 5 min*** washes with ***PBSTw***.

#### *Secondary probes hybridization*

6) For each column, to ***120 µL*** of ***PBSTw*** mix ***1–1.5 µL*** of ***10 µM*** secondary probe oligonucleotide complementary to the used SABER concatemer sequences for the primary probes *(Table S1)*.

- **Notes:** *when doing multiplex SABER TSA or combined AP SABER + SABER TSA, make a mix of multiple haptenized secondary probes.*

  *If doing SABER HCR or direct SABER FISH, add a desired mix of HCR adapter secondary initiator probes or fluorophore-labeled secondary probes, respectively.*

7) Pre-heat the secondary probe hybridization mix to **42°C**. Exchange ***PBSTw*** to ***120 µL*** of the mix. Leave hybridizing for ***1 h***.

8) Do ***1x*** buffer exchange and ***3x 5 min*** washes with ***PBSTw***.

9) Transfer the columns to room temperature.

- **Notes:** *pause point. If necessary, samples can be stored for **1-2 days** at **4°C**.*

  *If doing the direct SABER FISH protocol, proceed immediately to mounting of the samples (Section S4.5).*

***Anti-hapten antibody blocking and incubation (only for AP SABER or SABER TSA)***

10) Depending on the signal detection protocol used, choose the following blocking buffers and blocking times:

| ***Antibody type*** | ***Detection method*** | ***Blocking buffer*** |
|---|---|---|
| AP (Alkaline Phosphatase) conjugates | Colorimetric signal development with Vector Blue, NBT/BCIP, FastBlue, or FastRed | No blocking is necessary, do extra wash in ***PBSTw*** and incubate antibodies in plane ***PBSTw*** |
| POD/HRP (Horseradish peroxidase) conjugates | Tyramide Signal Amplification (TSA) FISH | Block for ***30 min*** in ***5% horse serum*** and ***5%*** stock ***Roche Western Blocking Reagent*** in ***PBSTw*** |

Exchange ***PBSTw*** for ***400 µL*** of an appropriate blocking buffer. Pre-heat the oven to **37°C**.

11) Prepare ***1:2000*** antibody dilutions in the corresponding blocking buffer for ***anti-DIG/FITC-AP*** or ***anti-DIG/FITC-POD*** antibodies. Incubate the samples in ***250 µL*** of the antibody solution for ***45 min*** at **37°C**. Put ***PBSTw*** at **37°C** for subsequent washes.

12) Still at **37°C**, gently flush out most of the antibodies, do ***1x*** buffer exchange to ***PBSTw*** followed by ***1x 5 min***, ***10 min***, ***15 min***, and ***30 min*** PBSTw washes.

Exchange ***PBSTw*** one extra time and either leave the samples overnight to wash or immediately proceed to signal development.

- **Note:** *pause point. If necessary, samples can be stored for **1-2 days** at **4°C**.*

13) Proceed to "<u>Day 3. Post-hybridization washes, 2nd hybridization, and development</u>" section of the protocol.

***SABER HCR: preparation of the hairpins and signal amplification***

10) Do ***1x*** buffer exchange and ***2x 10 min*** incubation in ***400 µL*** of ***HCR amplification buffer*** (***5xSSCTw***).

- **Note:** *if you have a saved previously used **HCR hairpin amplification mix**, proceed immediately to step (13).*

11) In separate PCR tubes, add ***2 µL*** of corresponding H1 and H2 ***3 µM*** hairpin stocks (*do not mix them together before they are annealed!*). Heat the tubes in a PCR thermocycler at **95°C** (**105°C** lid) for ***90 sec*** and cool to room temperature in a dark drawer for ***30 min***.

12) Mix all the pre-annealed HCR hairpins in ***200 µL*** of ***HCR amplification buffer***.

13) Drain off all the buffer from the columns and quickly add ***100-200 µL*** of the prepared ***HCR hairpin amplification mix*** to the columns. Leave the samples in a dark drawer at room temperature overnight (***16+ hours***).

### S4.4 Day 3. Signal development and sample mounting

*See the S3 Recipes section for the preparation of corresponding detection/development buffers for each substrate. Depending on a chosen signal development method proceed as follows:*

***Colorimetric signal development with AP substrates (Vector Blue, NBT/BCIP, FastBlue, or FastRed)***

1) Do ***1x*** buffer exchange and ***1x 5 min*** incubation in ***400 µL*** of the ***detection buffer*** (without the detection substrate).

2) Column-by-column, transfer worms from columns to a small petri-dish filled with the ***detection buffer*** to remove any accumulated dust, damaged animals and count worms. Then, transfer the worms into a well of a 24-well plate pre-filled with ***300 µL*** of the ***detection buffer*** (without substrate).

The following steps are preferably performed in a room without direct bright white light

3) Prepare fresh ***AP color development solution*** (detection buffer + substrate). Remove as much as possible of the detection buffer from the wells, trying to keep the worms in the inside rim of the wells. Add at least ***400 µL*** of the ***color development solution*** and gently swirl the plate to mix the worms with the solution. Start your timer and monitor the development under a stereomicroscope with bottom light on at first every ***5-10 min*** for strongly expressed genes or ***20-30 min*** for weaker expressed genes at room temperature.

- **Note:** *for some substrates (like* ***NBT/BCIP****) and weakly expressed genes development at* ***37°C*** *can be beneficial to shorten the development time. If the color development buffer noticeably changes color or the development takes longer than* ***3 hours****, prepare fresh color development buffer and transfer worms into it.*

4) To stop the development reaction, transfer worms in as little volume as possible to another well of the same 24-well plate pre-filled with ***1 mL*** of ***Mili-Q water*** with ***0.01% Tween 20***. Mix the worms in the well by pipetting to ensure efficient buffer exchange. Aspirate most of the ***Mili-Q-Tw*** and add another ***1 mL*** of ***Mili-Q-Tw***.

- **Note:** *for* ***NBT/BCIP*** *staining, a* ***5-10 min*** *wash in* ***96-100% EtOH*** *followed by another wash in* ***Mili-Q-Tw*** *can be beneficial for signal clarity and intensity before mounting (the color will change to blue from purple).*

5) For simple single-gene colorimetric staining, transfer the worms into another well of the same 24-well plate pre-filled with ***1 mL*** of the glycerol mounting solution.

- **Note:** *pause point. If necessary, samples can be stored for at least* ***1 month*** *at* ***4°C*** *before mounting on slides. If you want to proceed with mounting on the same day, incubate the worms for at least* ***1 h*** *at room temperature in* ***glycerol mounting solution****.*

***Fluorescent signal development by tyramide signal amplification (TSA)***

1) Prepare fresh ***TSA development buffer*** with a desired fluorescent tyramide. Incubate in ***250 µL*** of the ***TSA development buffe***r in the dark for ***10 min*** at room temperature.

- **Note:** *use **1/1000** dilution of **CF568 (red) tyramide (5 mM)**, **1/500** dilution of **FITC tyramide (5 mM)**, and **1/250** dilution of **CF647 (5 mM)**.*

2) Flush the ***TSA development buffer***, do ***2x*** brief buffer exchanges in ***PBSTw*** and then ***4x 5 min*** washes in ***PBSTw***.

- **Note:** *at this point you can take out some worms from the columns and check the signal and background under a fluorescent stereomicroscope. Continue washing for an extra **30 min** to destain the worms further.*

3) In case of a single gene development or the last step of the multiplex TSA protocol, add ***DAPI*** and incubate overnight at **4°C**.

4) Flush ***DAPI***, wash once for ***5 min*** with ***PBSTw***. Column-by-column, transfer worms from columns to a small petri-dish filled with ***PBSTw*** to remove any accumulated dust, damaged animals and count worms. Then, transfer the worms into wells of a 24-well plate pre-filled with ***1 mL*** of the ***glycerol mounting solution***.

- **Note:** *pause point. If necessary, samples can be stored for several days/weeks at **4°C** before mounting on slides. If you want to proceed with mounting on the same day, incubate the worms for at least **1 h** at room temperature in **glycerol mounting solution**.*

In case of continuing with the double/triple multiplex TSA FISH protocol, do ***1x*** buffer exchange and incubate ***1x 30 min*** with ***Sodium azide stop solution*** in the dark to inactivate a residual peroxidase activity. Wash ***4x 5 min*** in ***PBSTw***. Repeat all the steps starting from **Day 2: Anti-hapten antibody blocking and incubation** section with a different antibody.

***SABER HCR. Post-amplification washes and mounting***

1) Drain/collect as much as possible of the ***HCR hairpin amplification mix*** from the columns in a separate 1.5-2 mL tube. Keep saved HCR mix at ***-20ºC***.

- **Note:** ***HCR hairpin amplification mix*** can be reused at least 3 times without noticeable loss of sensitivity and signal brightness. Do not worry about sample drying when collecting the mix if it is done within 1-2 minutes.

2) Do ***1x*** buffer exchange and ***4x 5 min*** washes in ***400 µL*** of ***2xSSCTw***.

3) Still in columns, add ***DAPI*** and incubate overnight at **4ºC**.

4) Flush ***DAPI***, wash once for ***5 min*** with ***2xSSCTw***. Column-by-column, transfer worms from columns to a small petri-dish filled with ***2xSSCTw*** to remove any accumulated dust, damaged animals and count worms. Then, transfer the worms into wells of a 24-well plate pre-filled with ***1 mL*** of the ***glycerol mounting solution***.

- **Note:** *pause point. If necessary, samples can be stored for several days/weeks at **4ºC** before mounting on slides. Kepp in mind that some fluorophores bleach-out faster than others during storage. If you want to proceed with mounting on the same day, incubate the worms for at least **1 h** at room temperature in **glycerol mounting solution**.*

### ***S4.5 Mounting on slides***

*All steps are done with the assistance of a stereomicroscope. The same steps are applied when mounting in another medium, like VECTASHIELD.*

1) Move 2-4 worms stored in ***glycerol mounting solution*** in a small drop in the center of an ethanol-cleaned glass slide.

- **Note:** *it is hard to control orientation of more worms due to their tendency to roll on the sides during mounting.*

2) Using a plastic pipette tip, pick some Vaseline grease and apply it to the corners of a rectangle corresponding to the size of your coverslips to form supporting legs.

- **Note:** *Vaseline legs serve as a support for the coverslip weight, preventing it to break/overs squash the animals.*

3) Gently put an ethanol-cleaned coverslip on top of the Vaseline legs and gently, corner by corner, press it until the desired worm positioning is achieved.

- **Note:** *by controlling the pressure and height of the Vaseline legs, you should be able to gently slide the coverslip to achieve the best orientation of the worms.*

4) If the results are satisfactory, you can secure the borders of the slide by nail polish to prevent coverslip movement and sample drying.

### References

**Irvine, S. Q.** (2007). Whole-mount *in situ* hybridization of small invertebrate embryos using laboratory mini-columns. *BioTechniques* **43**, 764–768.

**Kishi, J. Y., Lapan, S. W., Beliveau, B. J., West, E. R., Zhu, A., Sasaki, H. M., Saka, S. K., Wang, Y., Cepko, C. L. and Yin, P.** (2019). SABER amplifies FISH: enhanced multiplexed imaging of RNA and DNA in cells and tissues. *Nat. Methods* **16**, 533–544.

**Lauter, G., Söll, I. and Hauptmann, G.** (2011). Two-color fluorescent *in situ* hybridization in the embryonic zebrafish brain using differential detection systems. *BMC Dev. Biol.* **11**, 43.
